# Resolving the estrogen paradox in hereditary retinal degeneration: Esr1 activation suppresses Tnf-α signaling as a photoreceptor self-protection mechanism

**DOI:** 10.1101/2025.09.07.674682

**Authors:** Yuting Li, Yadi Li, Jiarui Luo, Lan Wang, Qianlu Yang, Qianxi Yang, Cong Duan, Wenrong Xu, Yujie Dong, Lei Kong, Yan Li, Wenjia Zhang, Kangwei Jiao, Zhijian Zhao, Christina Schwarz, François Paquet-Durand, Junchuan Ye, Zhulin Hu, Jie Yan

**Author notes:** Yuting Li, Yadi Li and Jiarui Luo are shared first authors who contributed equally to this work. Jie Yan, Zhulin Hu, and Junchuan Ye are shared corresponding authors who contributed equally to this work. Correspondence (J.Y.); (ZL.H.); (JC.Y.).

## Abstract

Retinitis pigmentosa (RP) is an inherited retinal degenerative disorder characterized by progressive photoreceptor loss and irreversible blindness. Increasing evidence implicates neuroinflammation as a contributor to photoreceptor degeneration extending beyond the initial genetic insult. Although estrogen has been reported to exert anti-inflammatory effects in the central nervous system, its role in RP remains controversial, with some studies suggesting a paradoxical exacerbation of retinal pathology. To address this discrepancy, we identify estrogen receptor alpha (Esr1) as a central immunoregulatory hub in RP. Transcriptomic analyses of *rd1* and *rd10* revealed upregulation of estrogen-responsive and inflammatory pathways, with Esr1 expression markedly elevated during degeneration. TUNEL assays demonstrated that systemic estradiol (E2) exerted divergent effects, protective in *rd1* yet deleterious in *rd10,* whereas selective pharmacological activation of Esr1 with propyl pyrazole triol (PPT) consistently reduced photoreceptor death, preserved dark-adapted ERG responses, and downregulated inflammatory mediators including Tnf-α, Cx3cl1/Cx3cr1, Cd68, and Iba1. Mechanistically, Esr1 activation repressed microglial *Tnf* transcription and disrupted a self-sustaining Cx3cl1/Cx3cr1–Tnf-α signaling loop driving microglial recruitment, activation and neurotoxicity in the outer nuclear layer (ONL). Targeted interventions confirmed tumor necrosis factor receptor 1 (Tnfr1) as the principal mediator of Tnf-induced photoreceptor death: selective inhibition with R7050 conferred superior protection compared with broad-spectrum Tnf-α inhibitors (etanercept, infliximab). Cx3cr1 blockade likewise suppressed microglial activation and improved visual outcomes. Collectively, our findings establish Esr1 activation as not merely an external intervention but the amplification of an intrinsic self-protective program, positioning Esr1, Tnfr1, and Cx3cr1 as actionable therapeutic targets to suppress neuroinflammation and preserve vision in RP and related retinal disorders.

## Introduction

Hereditary retinal degenerations (IRDs) encompass a spectrum of genetic disorders that drive progressive photoreceptor loss and ultimately blindness [1]. Retinitis pigmentosa (RP), the most prevalent and currently untreatable IRDs, affects over two million individuals worldwide (≈1 in 4,000) [1]. RP typically presents with nyctalopia due to primary rod photoreceptor degeneration, followed by concentric constriction of the visual field (“tunnel vision”) driven by secondary cone degeneration, a consequence of rod death [1]. In the absence of disease-modifying therapies, delineating novel molecular targets is critical for halting or delaying RP progression.

The *rd1* and *rd10* mouse strains represent well-established RP models, each harboring *Pde6b* mutations encoding the β-subunit of rod cyclic guanosine monophosphate (cGMP) phosphodiesterase (PDE6β) [2]. These loss-of-function mutations impair cGMP hydrolysis, resulting in pathological cGMP accumulation [3], aberrant Ca² influx [4], and activation of non-apoptotic photoreceptor death pathways, notably Parthanatos [5]. Degeneration in *rd1* begins at postnatal day 8 (P8) and peaks around P13 [6], whereas *rd10* exhibits a delayed and more protracted course, with onset at P16-P20 and a peak at P21-P25 [7]. Pharmacological PDE6 inhibition by zaprinast mimics *Pde6b*-driven pathology by selectively blocking cGMP hydrolysis [8], thereby serving as a chemical surrogate for modeling cGMP-induced photoreceptor death [9]. However, interventions targeting cGMP accumulation [10], Ca² overload [4], or Parthanatos [11] have proven insufficient to fully halt photoreceptor degeneration, suggesting the existence of additional, yet unidentified pathogenic pathways and therapeutic targets.

Among these, microglia-driven neuroinflammation has emerged as a non-cell-autonomous contributor to retinal degeneration [12, 13]. Retinal microglia, the primary immune sentinels, transition from a homeostatic surveillant state to an activated phenotype, migrating to degenerating photoreceptors and releasing pro-inflammatory cytokines such as tumor necrosis factor alpha (TNF-α) [13, 14]. TNF-α engages its receptor TNFR1 to initiate neuroinflammatory signaling [15], and the resulting sustained inflammatory milieu further accelerates neuronal loss [13, 14]. Microglial homeostasis is tightly regulated by chemokine (C-X3-C motif) ligand 1 (CX3CL1), and its receptor CX3CR1, where neuron- and photoreceptor-derived CX3CL1 maintains microglial quiescence [16, 17]. Disruption of this axis during degeneration likely amplifies microglial activation and exacerbates photoreceptor death [17–19].

Given the role of neuroinflammation in retinal degeneration, hormonal modulation has attracted attention as a potential strategy to restrain microglia-mediated damage [20]. Estrogen, particularly 17β-estradiol (E2), is well recognized for its neuroprotective and anti-inflammatory properties within the central nervous system [20, 21]. In the retina, estrogen has been proposed as an endogenous self-protective mechanism engaged in response to stress [22]; however, the precise pathways by which it confers such protection remain poorly defined. Among the four natural isoforms (estrone, estradiol, estriol, estetrol), E2 is the most potent and biologically active, acting primarily through estrogen receptor α (ERα/ESR1), with additional contributions from ERβ (ESR2) and the G-protein-coupled estrogen receptor (GPER) [20, 23]. Despite this protective reputation, accumulating evidence indicates that estrogen’s actions in neurodegeneration are context-dependent, with some studies reporting exacerbation of pathology under certain conditions [24, 25], raising controversy over its therapeutic value in IRDs.

To resolve this controversy, we integrated bulk and single-cell transcriptomic analyses across multiple *Pde6b*-associated retinal degeneration models. Bulk RNA-seq of *rd1* and *rd10* retinas revealed progressive activation of estrogen-responsive and inflammatory pathways, along with upregulation of microglial markers including *Cx3cl1*, *Cx3cr1*, *Cd68* and *Aif1*. Single-cell profiling confirmed microglial expansion, localized *Tnf* expression, and loss of estrogen-responsive gene signatures in mutant photoreceptor that were otherwise enriched in *wt* counterparts. CellChat analysis further identified disease-specific microglia–rod interactions that were absent in *wt* but established in *Pde6b* mutants. Immunostaining demonstrated markedly enhanced Esr1 expression in degenerating retinas. Functionally, while E2 conferred neuroprotection in *rd1*, it aggravated degeneration in *rd10*, highlighting genotype-dependent effects. In contrast, selective Esr1 activation uniformly attenuated photoreceptor loss in both models by suppressing *Tnf* and downstream inflammatory cascades. Blockade of Tnfr1 provided more robust neuroprotection than pan-Tnf-α inhibition, and microglia-specific Cx3cr1 inhibition confirmed microglia as the principal source of Tnf-α. *In vivo*, intravitreal administration of Esr1 agonists, Tnf-α antagonists, or Cx3cr1 inhibitors preserved retinal structure and function in both genetic and chemically induced degeneration models. Collectively, these findings position Esr1 as an intrinsic protective regulator in photoreceptor degeneration and identify selective Esr1 activation as a potent, inflammation-targeted, genotype-independent therapeutic strategy that surpasses systemic estrogen in the treatment of inherited retinal diseases.

## Materials and methods

### Animals

Experiments involved C3H *Pde6b^rd1/rd1^* (*rd1*), C57BL/6J *Pde6b^rd10/rd10^* (*rd10*), congenic C3H *Pde6b^+/+^* wild-type (C3H), and C57BL/6J *Pde6b^+/+^*wild-type (C57) mice. All procedures were under the supervision and assessment by the Laboratory Animal Welfare Ethics Committee Yunnan University. *rd1* and *rd10* were used to compared with their congenic wild-type animals (C3H and C57) respectively regardless of gender.

### Retinal explant culture

Retinal explants from *rd1*, C3H, *rd10*, and C57 mice were prepared as described [26]. Retinas were collected at P5 for *rd1* and C3H, and P12 for *rd10* and C57. The two retinas from each animal were divided across experimental groups to maximize independent observations. After 48h in culture, explants were treated with 10μM E2 [27] (HY-B0141; MedChemExpress, Sollentuna, Sweden), 10μM E2 plus 10nM AZD9496 [28] (HY-12870; MedChemExpress), 15nM PPT [29] (HY-100689; MedChemExpress), 0.63nM Etanercept [30] (ETN, HY-108847; MedChemExpress), 6.7μM Infliximab [31] (IFX, HY-P9970; MedChemExpress), 65μM R7050 [32] (HY-110203; MedChemExpress), or 4μM AZD8797 [33] (HY-13848; MedChemExpress). Compound concentrations were based on retinal dose-response curves (Figure S1A-G). Cultures were terminated at P11 (*rd1*) or P20 (*rd10*) by fixation with 4% paraformaldehyde. Explants were embedded in Tissue-Tek (Sakura Finetek, Alphen aan den Rijn, The Netherlands) and cryosectioned (12μm) using a CryoStar NX50 (Thermo Fisher Scientific, Runcorn, UK).

### Intravitreal injections

Single intravitreal injections were performed in *rd10* and C57 mice from P16 to P23 as described [34]. Mice were anesthetized by intraperitoneal tribromoethanol (83.3 mg/kg). In *rd10* mice, the left eye received 0.1μL of PSB mixed with DMSO containing 1.5μM PPT, 31.5nM ETN, 335μM IFX, 3.25mM R7050, or 200μM AZD8797, while the right eye received 0.1μL of an equal volume of PBS mixed DMSO as a sham control. In C57 mice, 20mM zaprinast [9] (ZAP, HY-B1816; MedChemExpress) was co-administered with each compound in the left eye (0.1μL, 400μM final intraocular concentration, selected based on Figure S1H), with the right eye again receiving a DMSO sham injection. Final intraocular concentrations were calculated based on a 5μL mouse ocular volume [34, 35], matching those used in retinal explant cultures.

### Dark-adapted electroretinography (ERG)

Dark-adapted ERGs were performed using the Celeris system (Diagnosys LLC, Lowell, MA, USA). Intravitreal injections were administered in *rd10* and C57BL/6J mice from P16 to P21, coinciding with the critical window of functional decline before near-complete loss of retinal responses at ∼P23 [36, 37]. Histological analyses were extended to P23 to capture cumulative structural degeneration beyond the functional floor effect [36, 37]. For all experiments, *rd10* and ZAP-treated C57 mice served as paired controls, with the right eye as control and the left as experimental.

Mice were dark-adapted for ≥12h before testing. Anesthesia was induced by intraperitoneal injection of tribromoethanol (83.3mg/kg), and pupils were dilated with Tropicamide-Phenylephrine Eye Drops (Santen Pharmaceutical Co., Ltd., Osaka, Japan). dark-adapted responses were elicited by 3cd·s/m² flash stimuli, and signals were recorded within a 300ms window. A-wave amplitudes were measured from baseline to the negative trough, and b-wave amplitudes from the a-wave trough to the subsequent positive peak. For comparative consistency, all a-wave values were converted to absolute values, thereby representing negative amplitudes as positive numbers. All recordings were performed under dim red light, and data acquisition and analysis were conducted using the integrated Celeris platform.

### Single-cell RNA-seq (scRNA-seq) data processing and analysis

Raw sequencing data (eyeballs collected from P19 *rd10* and C57) were processed using Cell Ranger (v9.0.0) with the Mus musculus GRCm39 reference genome, and UMI count matrices were analyzed in Seurat (v5.0.3). Cells with <200 detected genes, <1000 UMIs, log10 (Genes/UMI) <0.7, >30% mitochondrial gene content, and >5% hemoglobin gene content were excluded. Doublets were removed using DoubletFinder. Data were normalized, integrated using Harmony for batch correction, scaled, and clustered using PCA and the Louvain algorithm, and visualized by t-SNE. Cell types were annotated with canonical markers. UCell scoring was applied to assess pathway activity, and t-SNE plots were generated to visualize differential expression across conditions. Cell-cell communication analysis was performed using CellChat (v1.6.1) with the mouse ligand-receptor database.

### Bulk RNA sequencing (RNA-seq)

Total RNA was isolated from pooled *rd1*, *rd10* and congenic *wt* mouse eyeballs (both eyes per animal) or from PPT-treated *rd1* and *wt* retinal explants. Total RNA was extracted using standard protocols and quality-checked with the RNA Nano 6000 Assay Kit (Agilent Technologies, USA) and NanoPhotometer® spectrophotometer (IMPLEN, USA). Sequencing libraries were constructed using the NEBNext® Ultra™ RNA Library Prep Kit for Illumina® (NEB, USA) and sequenced on an Illumina NovaSeq platform to generate 150bp paired-end reads. The mRNA expression comparison between *rd1* and *wt* from P7 to P21 employed datasets downloaded from the GSE62020 [38].

### GSEA algorithm

GSEA was performed using GSEA software (v.4.3.3) in conjunction with the Molecular Signatures Database (v.7.4). One thousand permutations were used.

### Hub-Gene identification

Differential genes between *rd1/rd10* and congenic *wt* retinas were called in R 4.5.1 (Bioconductor) with cut-offs fold change (FC) > 1.5 and *p* < 0.05. The resulting protein–protein interaction network was retrieved from STRING and analysed in Cytoscape 3.10. Six algorithms (Degree, MNC, MCC, Closeness, Stress, EPC) ranked nodes; genes present in the top 10 list of every metric were designated as hubs.

### TUNEL staining

TUNEL staining was performed using a commercial kit (Beyotime, Jiangsu, China) to detect dying cells. Retinal explants and eye cup sections were dried, stored at −20 °C, rehydrated in 0.1M PBS, and permeabilized with 0.5% Triton X-100 for 5 min. Sections were washed in PBS and incubated with 100μL of TUNEL solution (enzyme:labeling solution, 1:9) in a light-protected humid chamber at 37 °C for 60 min. After additional PBS washes, sections were counterstained with DAPI (G1407; Servicebio, Wuhan, China) and imaged using confocal microscopy.

### Immunohistochemistry

Sections were rehydrated in PBS for 15 min and blocked with 10% normal goat serum, 1% bovine serum albumin, and 0.3% PBST for 1h. Primary antibodies: rabbit anti-Esr1 (1:200; YT1634; Immunoway, California, USA), rabbit anti-Cd68 (1:200; BA3638; BOSTER, California, USA), rabbit anti-Iba1 (1:400; ym8165; Immunoway), rabbit anti-Cx3cl1 (1:200; YT5354; Immunoway), rabbit anti-Cx3cr1 (1:200; YT5112; Immunoway), and rabbit anti-Tnf-α (1:200; YM8306; Immunoway), rabbit anti-Tnfr1(1:50; YT4687; Immunoway), Mouse anti-Glutamine Synthetase (1:1000; MAB302; Abcam)-were diluted in blocking solution and incubated overnight at 4°C. Sections were rinsed in PBS (3 × 10 min) and incubated with goat anti-rabbit Alexa Fluor 488 (1:350; A11034; Molecular Probes, Oregon, USA) for 1h. After an additional PBS rinse (3 × 10 min), sections were mounted with DAPI-containing mounting medium (Servicebio).

### Microscopy and image analysis

Retinal explant and *in vivo* samples were imaged using a Zeiss LSM 900 confocal microscope (Carl Zeiss, Oberkochen, Germany). Z-stack and tile scans were acquired at 20× magnification, with 12 µm sections captured across four optical planes. Images were processed using ZEN (Blue edition) software. Positive cells in the outer nuclear layers (ONL) were quantified by manually counting DAPI-stained nuclei in six rectangular regions to estimate mean cell density, which was used to calculate total ONL cell numbers. The percentage of positive cells was calculated as the ratio of positive to total cells, and signal intensities were quantified using Zeiss Zen 3.0 software (Carl Zeiss).

### Statistical analysis and software use

Two-way comparisons were analyzed using paired t-tests or student’s t-test. Multiple comparisons were assessed by one-way ANOVA with Tukey’s post hoc test. Analyses were performed using GraphPad Prism 8 (GraphPad Software, La Jolla, CA, USA), with *p* < 0.05 considered significant. Significance levels were denoted as follows: *p* < 0.05 (*), *p* < 0.01 (**), *p* < 0.001 (***), and *p* < 0.0001 (****). Data in Figures 7 and S1 were normalized by linear scaling χ scaled = (χ – χ min)/(χ max – χ min) using SPSS Statistics 26 (IBM, Armonk, NY, USA). Spearman correlation analyses were conducted in R (v4.0.1). Figures were generated using Photoshop 2025 and Illustrator 2025 (Adobe, San Jose, CA, USA), and Figure 8 was created with BioRender.com.

## Results

### Esr1 activation as an intrinsic self-protective mechanism to attenuate photoreceptor degeneration and preserves retinal function

To assess the involvement of estrogen signaling in *Pde6b*-associated photoreceptor degeneration, we performed gene set enrichment analysis (GSEA) on RNA-seq datasets from *rd1* (P13) and *rd10* (P21) retinas. Both models exhibited significant enrichment of estrogen-responsive pathways (Figure 1A, Figure S2A), implicating a potential role for estrogen signaling in disease progression. scRNA-seq profiling further revealed that estrogen response gene set were highly enriched in *wt* photoreceptor but markedly diminished in *rd10* counterpart, while microglia and Müller cells in *rd10* retained strong signatures (Figure 1B). Immunofluorescence staining corroborated this finding, showing robust upregulation of Esr1 protein throughout the degenerating retina, with predominant localization in the inner segment (IS), inner nuclear layer (INL), and ganglion cell layer (GCL) in both *rd1* and *rd10* models compared to *wt* controls (Figure 1C–F).

**Figure 1.**
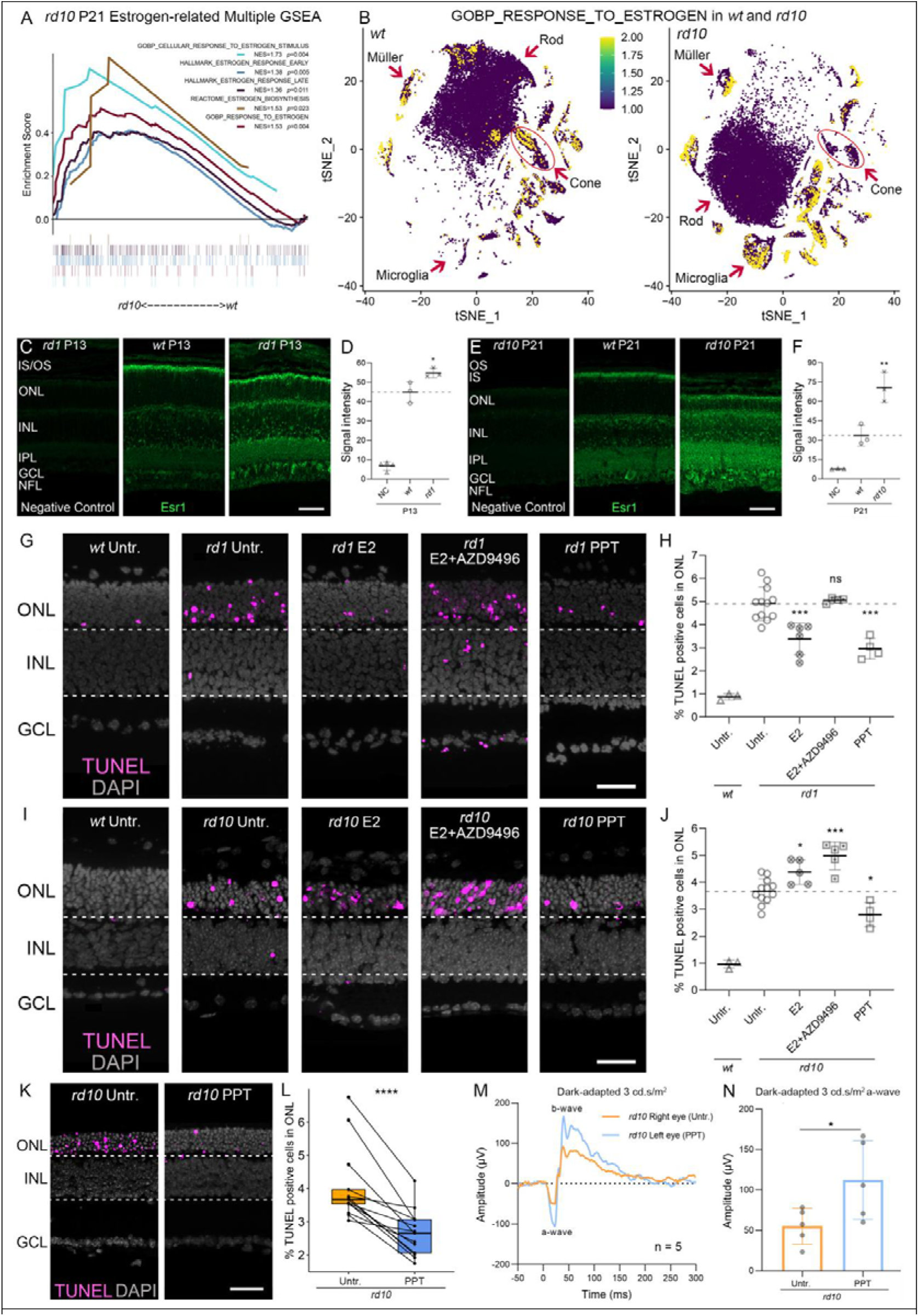
Esr1 as an intrinsic self-pretective mechanism to preserve photoreceptor viability and function. **(A)** GSEA of RNA-seq data from *rd10* retinas at postnatal day 21 (P21) revealed significant positive enrichment of Estrogen-related pathways. **(B)** t-SNE plots of UCell scores for the GOBP RESPONSE TO ESTROGEN gene set in *wt* and *rd10* retinal cells. Yellow indicates high scores, purple indicates low scores. Arrows highlight Müller cells, rods, cones and microglia. Notably, estrogen response gene set scores were enriched in *wt* photoreceptor but markedly reduced in *rd10* photoreceptor. Microglia retained high estrogen response signatures in *rd10*. **(C-D)** Immunofluorescence staining for Esr1 (green) demonstrated increased signal intensity in the entire retina of *rd1* mice compared to congenic *wt* controls and negative staining controls. **(E-F)** Esr1 (green) expression was also elevated in *rd10* (P21) retinas relative to *wt* and negative controls. Esr1 was predominantly localized to IS, INL, and GCL. **(G-J)** TUNEL staining (magenta) of ONL in *rd1* and *rd10* retinal explants versus wild type (*wt*). Explants were untreated (Untr.) or treated with 10μM E2, E2 + 10nM Esr1 antagonist AZD9496, or 15nM Esr1 agonist PPT. Scatter plots showed percentages of TUNEL-positive cells (H,J). **(K-N)** Intravitreal PPT in *rd10* reduced ONL TUNEL positivity (K,L) and improved dark-adapted ERG a-wave amplitudes (M,N). Error bars: SD; significance levels: * = *p* < 0.05; ** = *p* < 0.01; *** = *p* < 0.001; **** = *p* < 0.0001. OS = outer segment, IS = inner segment, ONL = outer nuclear layer, INL = inner nuclear layer, IPL = inner plexiform layer, GCL = ganglion cell layer, NFL = nerve fiber layer; scale bar = 50 µm. Dashed lines indicate untreated mutant or *wt* baseline levels. DAPI (grey) was used as nuclear counterstain.

To functionally probe estrogen signaling, we employed retinal explant cultures treated with 17β-estradiol (E2), E2 plus the Esr1 antagonist AZD9496, or the Esr1-selective agonist PPT. TUNEL assays confirmed robust outer nuclear layer (ONL) cell death in *rd1* and *rd10* explants relative to *wt* (Figure 1G-J). E2 administration significantly attenuated ONL cell death in *rd1* but, conversely, exacerbated degeneration in *rd10*. The addition of AZD9496 abrogated E2-mediated neuroprotection in *rd1* and further aggravated photoreceptor loss in *rd10*, supporting a genotype-dependent and context-specific effect of endogenous estrogen signaling. In contrast, selective Esr1 activation *via* propyl pyrazole triol (PPT) consistently reduced ONL TUNEL positivity in both models, underscoring the genotype-independent neuroprotective role of direct Esr1 stimulation (Figure 1G-J). Dose-response curves were generated to select appropriate treatment concentrations (Figures S1A-C).

*In vivo* intravitreal PPT administration in *rd10* mice, with contralateral eyes as controls, significantly reduced ONL TUNEL positivity (Figure 1K,L) and improved dark-adapted ERG responses, with prominent recovery of a-wave amplitudes (Figure 1M,N). Pharmacological induction of *Pde6b*-associated pathology by ZAP in *wt* mice increased ONL TUNEL positivity (Figure S2B,C) and suppressed ERG amplitudes relative to untreated eyes (Figure S2D,E). A dose-response curve for the effects of ZAP was compiled (Figure S1H) and a concentration of 400µM was chosen for further experiments. PPT co-administration attenuated ZAP-induced photoreceptor degeneration (Figure S2F,G) and restored ERG responses, particularly a-wave amplitudes (Figure S2H,I).

Together, these data indicate that targeted activation of Esr1, rather than non-specific estrogen stimulation, confers robust neuroprotection and preserves retinal function across both genetic and pharmacological models of retinal degeneration. These findings further support the notion that Esr1 signaling serves as an intrinsic, protective mechanism in the degenerating retina and represents a promising therapeutic target for retinal dystrophies.

### Esr1 activation reduces photoreceptor cell death *via* Tnf-**α** suppression

To delineate downstream pathways of Esr1, GSEA of *rd1* (P13) retinal RNA-seq revealed positive enrichment of Tnf-associated and inflammatory pathways (Figure S3A,E). Immunostaining confirmed elevated Tnf-α in *rd1* and *rd10* retinas versus *wt* and negative controls (Figure S3B-D), indicating Tnf-α–driven neuroinflammation contributes to degeneration. Following PPT treatment, GSEA demonstrated negative enrichment of Tnf-related and inflammatory pathways (Figure 2A, Figure S3F). CytoHubba analysis of PPT-treated *rd1* transcriptomes identified *Tnf* as a shared hub gene among six algorithms (Figure 2B), with significantly reduced *Tnf* expression. (Figure 2C).

**Figure 2.**
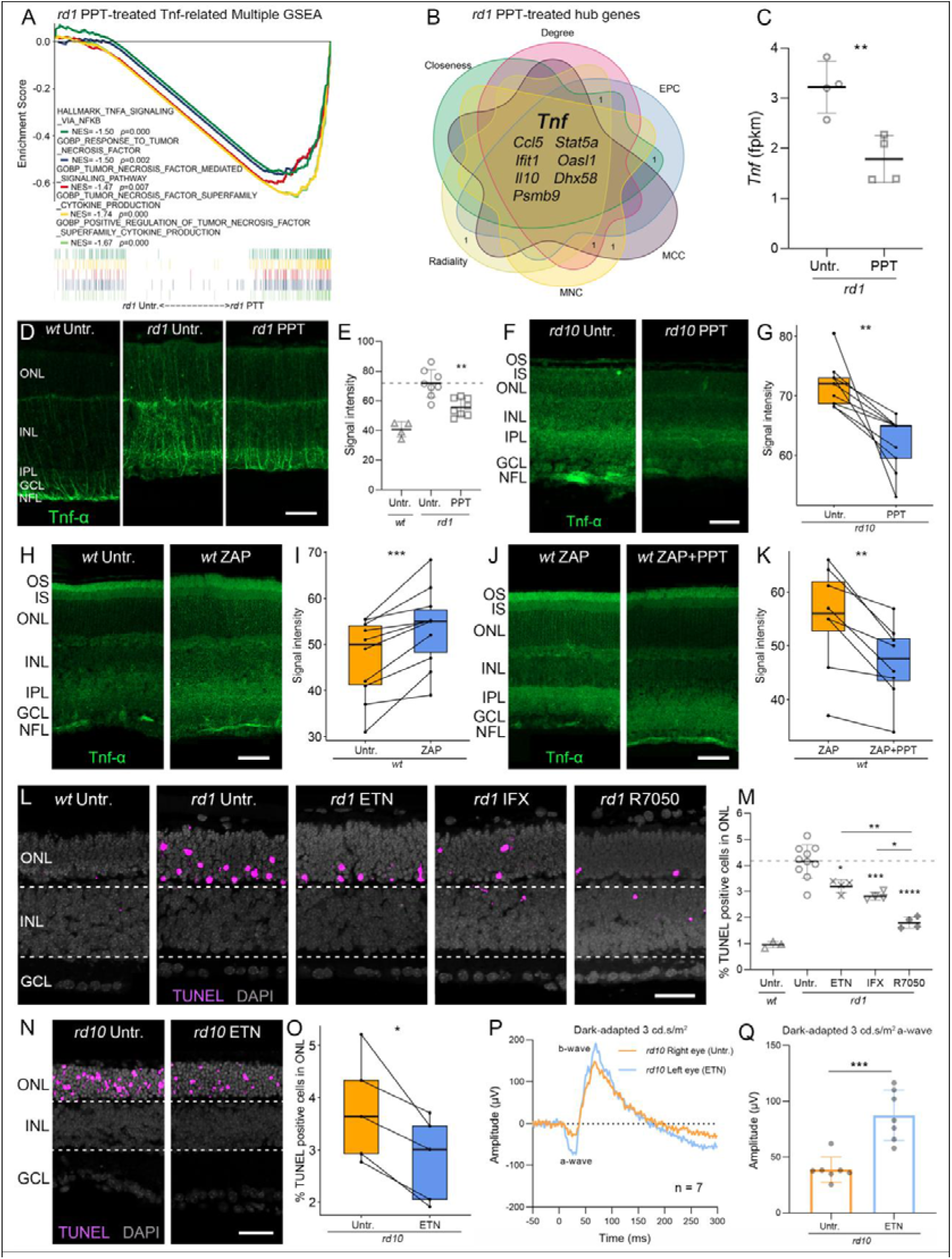
Esr1 activation suppresses Tnf-α signaling to protect photoreceptors. **(A)** GSEA of *rd1* retinal RNA-seq from explants showed negative enrichment of Tnf associated pathways after PPT treatment. **(B)** CytoHubba analysis identified *Tnf* as a common hub gene post-PPT. **(C)** *Tnf* expression (FPKM) is significantly reduced by PPT. **(D-K)** Immunostaining for Tnf-α (green) reveals elevated signal in *rd1* and *rd10* versus *wt*; PPT reduces Tnf-α in explant and *in vivo* models (D-G). ZAP elevated Tnf-α in *wt*, which was reversed by PPT (H-K). Notably, Tnf-α localized predominantly to Müller cells. **(L-M)** TUNEL assays (magenta) with DAPI (grey) as nuclear counterstain in *rd1* explants showed decreased ONL cell death with Tnf-α blockade (ETN, IFX) and further protection by the Tnf receptor antagonist R7050. **(N-Q)** Intravitreal ETN in *rd10* reduced ONL TUNEL positivity (N,O) and restored dark-adapted ERG a-wave amplitudes (P,Q). Error bars: SD; significance levels: * = *p* < 0.05; ** = *p* < 0.01; *** = *p* < 0.001; **** = *p* < 0.0001. OS = outer segment, IS = inner segment, ONL = outer nuclear layer, INL = inner nuclear layer, IPL = inner plexiform layer, GCL = ganglion cell layer, NFL = nerve fiber layer; scale bar = 50 µm. Dashed lines indicate untreated levels.

Consistently, immunostaining showed low Tnf-α signal in *wt*, robust elevation in *rd1* and *rd10* retina, particularly Müller cells, and significant reductions following PPT treatment in both explants and *in vivo* models (Figure 2D-G). In *wt*, ZAP markedly increased Tnf-α expression, while co-treatment with PPT reversed this effect (Figure 2H-K).

To verify the functional role of Tnf-α, *rd1* explant cultures were treated with the Tnf-α inhibitor ETN, the monoclonal antibody IFX, or the Tnf receptor antagonist R7050. Dose-response curves were generated to select appropriate treatment concentrations (Figures S1D-F). TUNEL assays demonstrated significantly fewer ONL dying cells in *wt* versus *rd1* (Figure 2L,M). ETN and IFX both reduced photoreceptor death, whereas R7050 yielded the most pronounced protection, surpassing untreated *rd1* and single-agent treatments (Figure 2L,M).

Taken together, these findings indicate that Esr1 activation mitigates photoreceptor degeneration in *Pde6b* mutant models by suppressing Tnf-α–mediated neuroinflammation.

### Inhibition of Tnf-**α**/Tnfr1 signaling protects photoreceptors and restores retinal function

To validate the pathogenic role of Tnf-α signaling *in vivo*, *rd10* and ZAP-induced *wt* mice were treated with either the soluble Tnf-α inhibitor etanercept (ETN), the monoclonal antibody infliximab (IFX), or the Tnfr1-selective antagonist R7050. In *rd10* mice, ETN treatment markedly reduced TUNEL-positive cells in ONL (Figure 2N,O) and significantly improved darked-adapted ERG responses, particularly a-wave amplitudes, indicative of photoreceptor functional preservation (Figure 2P,Q). Similarly, ETN attenuated ZAP-induced photoreceptor loss in *wt* mice and restored ERG amplitudes (Figure 3A-D).

**Figure 3.**
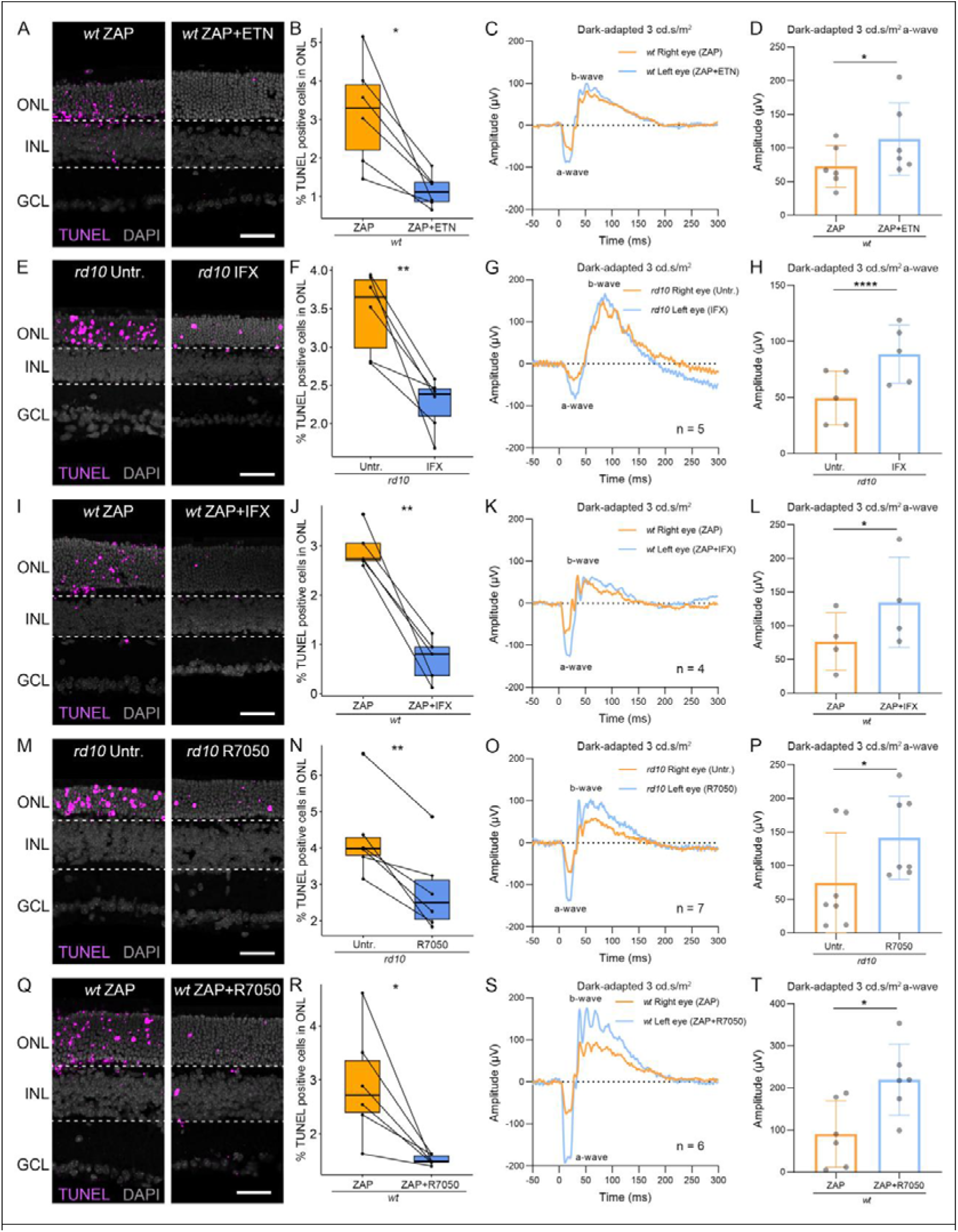
Tnf-α and Tnfr blockade protect photoreceptors and restore retinal function. **(A-D)** ETN reduced ZAP-induced ONL TUNEL positivity (A,B) and restored dark-adapted ERG responses (C) and a-wave amplitudes (D) in *wt*. **(E-H)** IFX lowered ONL TUNEL-positive cells in *rd10* (E,F) and improved ERG responses (G,H). **(I-L)** IFX also mitigated ZAP-induced ONL death (I,J) and rescued ERG function (K,L) in *wt*. **(M-P)** R7050 markedly decreased ONL TUNEL positivity in *rd10* (M,N) and enhanced ERG responses (O,P). **(Q-T)** R7050 further reduced ZAP-induced retinal degeneration in *wt* (Q,R) and restored dark-adapted ERG responses (S) and a-wave amplitudes (T). Error bars: SD; significance levels: * = *p* < 0.05; ** = *p* < 0.01;. ONL = outer nuclear layer, INL = inner nuclear layer, GCL = ganglion cell layer, scale bar = 50 µm; TUNEL assays (magenta) with DAPI (grey) as nuclear counterstain.

IFX exerted comparable neuroprotective effects. In *rd10* mice, IFX significantly reduced ONL cell death (Figure 3E,F) and enhanced ERG a-wave amplitudes (Figure 3G,H). In the ZAP-induced *wt* degeneration model, IFX preserved retinal structure and photoreceptor function, as evidenced by reduced TUNEL staining and improved a-wave responses (Figure 3I-L). However, IFX failed to restore b-wave amplitudes in both *rd10* and ZAP-treated *wt* mice (Figure 3G,K), suggesting incomplete recovery of inner retinal signaling.

Among the three agents, R7050 provided the most robust protection. R7050 profoundly suppressed photoreceptor apoptosis in *rd10* (Figure 3M,N), significantly restored both a- and b-wave ERG responses (Figure 3O,P), and completely reversed ZAP-induced retinal degeneration in *wt* mice (Figure 3Q-T).

scRNA-seq revealed minimal expression of *Tnfrsf1a* (Tnfr1) in rod photoreceptors of *wt* mice, whereas its expression was notably upregulated in *rd10* rods (Figure S4A). Temporal analysis further confirmed *Tnfrsf1a* upregulation during retinal degeneration in both *rd1* and *rd10* models (Figure S4B,C). Immunohistochemical analysis demonstrated Tnfr1 protein distribution throughout the retina, with prominent localization in Müller glia and photoreceptor inner segments (Figure S4D). Notably, overall Tnfr1 protein levels did not differ between *rd1*, *rd10*, and their respective congenic *wt* controls (Figure S4E,F).

Collectively, these *in vivo* findings, supported by transcriptomic and immunohistochemical data-identify Tnf-α/Tnfr1 signaling as a central driver of photoreceptor degeneration in *Pde6b*-associated retinal disease. Pharmacological disruption of this axis, particularly *via* selective Tnfr1 inhibition, effectively preserves retinal structure and visual function, offering a promising therapeutic strategy for inherited retinal degenerations.

### Microglia-derived Tnf-**α** as a putative driver of photoreceptor degeneration

To identify the cellular source of Tnf-α in degenerating retinas, we performed scRNA-seq in *wt* and *rd10* mice. Compared to *wt* retinas, the *rd10* retina exhibited a substantial expansion of the microglial population (Figure 4A). Transcriptomic mapping revealed that *Tnf* expression was predominantly restricted to microglia, with minimal expression detected in other retinal cell types (Figure 4B). Cell-cell communication analysis using CellChat further demonstrated a lack of microglia-photoreceptor interactions in *wt* mice (Figure 4C), whereas such crosstalk was prominently established in the degenerating *rd10* retina (Figure 4D), indicating disease-specific engagement.

**Figure 4.**
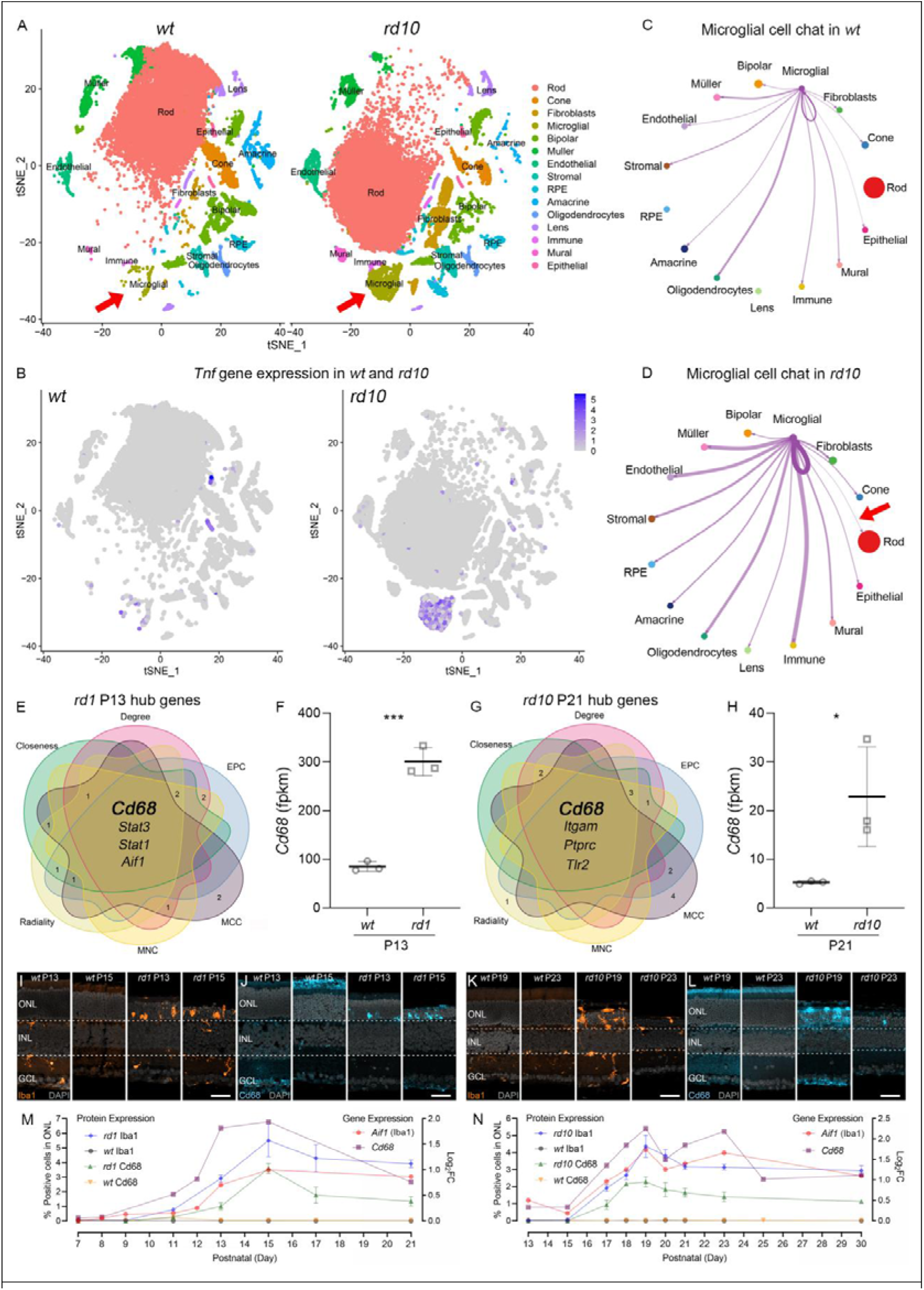
Microglia-derived Tnf-α mediates photoreceptor degeneration in *Pde6b*-mutant retinas. **(A)** Single-cell RNA sequencing (scRNA-seq) of whole eyeballs from *rd10* and congenic wild-type (*wt*) mice revealed an expanded microglial population in *rd10* retinas. **(B)** Feature plot showing *Tnf* transcripts predominantly localized to microglia rather than other retinal cell types. **(C,D)** Cell-cell communication analysis (CellChat) demonstrating absence of microglia-rod photoreceptor interactions in *wt* retinas (C), but emergence of such interactions in the *Pde6b*-mutant context (D). **(E-H)** CytoHubba analysis of *rd1* and *rd10* transcriptomes identified *Cd68* as a shared hub gene, with elevated gene expressions, indicating microglial overactivation. **(E-N)** Immunostaining and gene expression analyses showed a time-dependent upregulation of Iba1 (orange, *Aif1*) and Cd68 (cyan) during degeneration in *rd1* (P9-P21) and *rd10* (P13-P30) mice, relative to age-matched *wt* controls. Protein expression is plotted against the left Y-axis, while gene expression (lines) corresponds to the right Y-axis in dual-axis graphs.Error bars: SD; significance levels: * = *p* < 0.05; *** = *p* < 0.001. ONL = outer nuclear layer, INL = inner nuclear layer, GCL = ganglion cell layer; scale bar = 50µm. DAPI (grey) was used as nuclear counterstain.

To elucidate microglial drivers of degeneration, we next applied CytoHubba analysis across transcriptomic datasets from *rd1* and *rd10* retinas at their respective peak phases of photoreceptor loss. *Cd68* emerged as a convergent hub gene in both models, ranking top across six centrality algorithms and exhibiting significant upregulation in *rd1* and *rd10* mice (Figure 4E-H). This aligned with robust microglial activation observed histologically and transcriptionally. Immunostaining and RNA-seq confirmed progressive increases in Iba1 (*Aif1*) and Cd68 expression from P9 to P21 in *rd1*, and from P13 to P30 in *rd10*, relative to their *wt* counterparts (Figure 4I-N; Figure S5A-D; Figure S6A-D).

Collectively, these findings highlight microglia as the principal source of Tnf-α in the degenerating retina and implicate microglia-photoreceptor interactions as a possible pathological axis in *Pde6b*-associated retinal degeneration.

### Cx3cr1/Cx3cl1 signaling drives microglial activation and photoreceptor degeneration

Among chemokine pathways, the Cx3cl1-Cx3cr1 axis emerged as a central regulator of microglial activation during retinal degeneration [39]. In both *rd1* and *rd10* models of RP, transcript and protein levels of Cx3cl1 and its receptor Cx3cr1 were significantly upregulated during the critical windows of photoreceptor loss (Figure 5A-F; Figure S7A-D; Figure S8A-D), implicating this pathway in disease progression.

**Figure 5.**
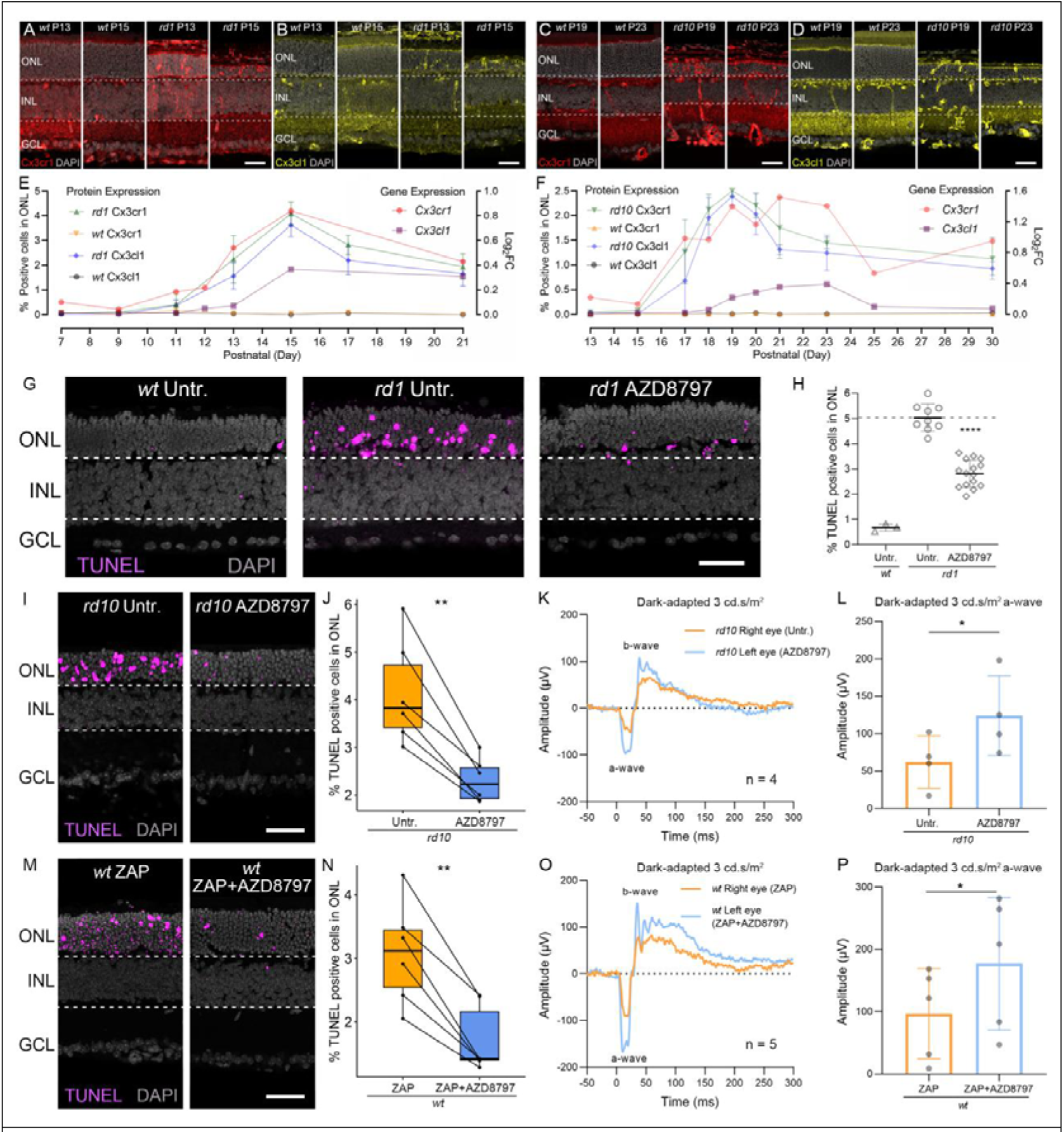
Cx3cl1-Cx3cr1 signaling mediates microglia-driven photoreceptor degeneration in *Pde6b*-mutant models. **(A-F)** Immunostaining and gene expression analyses showed a time-dependent upregulation of Cx3cl1 (yellow) and Cx3cr1 (red) during degeneration in *rd1* (P9–P21) and *rd10* (P13–P30) mice, relative to age-matched *wt* controls. Protein expression is plotted against the left Y-axis, while gene expression (lines) corresponds to the right Y-axis in dual-axis graphs.**(G-H)** Cx3cr1 inhibitor AZD8797 reduced ONL TUNEL-positive cells in *rd1* explants. Dashed lines indicate untreated levels. **(I-L)** In *rd10 in vivo*, AZD8797 decreased ONL cell death, restored dark-adapted ERG responses, and enhanced a-wave amplitudes. **(M-P)** AZD8797 mitigated ZAP-induced photoreceptor degeneration in *wt*, improving ERG function and a-wave amplitudes. * = *p* < 0.05; ** = *p* < 0.01; **** = *p* < 0.0001. ONL = outer nuclear layer, INL = inner nuclear layer, GCL = ganglion cell layer, scale bar = 50 µm; TUNEL assays (magenta) with DAPI (grey) as nuclear counterstain.

To assess whether microglial activation *via* this axis causally contributes to photoreceptor degeneration, we administered the Cx3cr1 antagonist AZD8797 in three complementary models: *rd1* retinal explants, *in vivo rd10* mice, and ZAP-induced photoreceptor injury in *wt* mice. In all contexts, AZD8797 treatment led to a significant reduction in ONL TUNEL-positive cells (Figure 5G-J, M-N), indicating attenuated photoreceptor cell death. Functional assessment *via* dark-adapted ERG revealed that AZD8797 robustly restored retinal responses, particularly improving a-wave amplitudes that reflect photoreceptor integrity (Figure 5K-L, O-P). A dose-response curve established 4µM as the optimal concentration for *ex vivo rd1* explants (Figure S1G).

Together, these results establish Cx3cl1-Cx3cr1 signaling as a key mediator of microglia-driven photoreceptor degeneration in *Pde6b*-associated RP. Pharmacological blockade of this axis using AZD8797 confers significant neuroprotection, preserving both retinal structure and visual function across multiple degenerative contexts.

### Targeting Esr1, Tnf-**α** signaling, and Cx3cr1 suppresses microglial recruitment, Cx3cl1/Cx3cr1 signaling, and neuroinflammation

We next examined the impact of Esr1 activation, Tnf-α inhibition, and Cx3cr1 blockade on microglial recruitment and neuroinflammatory signaling across *rd1* explants, *rd10* retinas, and ZAP-induced *wt* degeneration. Treatment with the Esr1 agonist PPT, Tnf-α inhibitors (ETN, IFX), the Tnfr1 antagonist R7050, or the Cx3cr1 inhibitor AZD8797 markedly suppressed microglial recruitment, as evidenced by reduced Iba1 expression within the ONL in all models (Figure 6A,B; Figure S9A,E; Figure S10A,B). Cd68 expression followed a similar trend: minimal in untreated *wt*, robustly induced during degeneration, and significantly attenuated upon intervention (Figure 6C,D; Figure S9B,F; Figure S10C,D).

**Figure 6.**
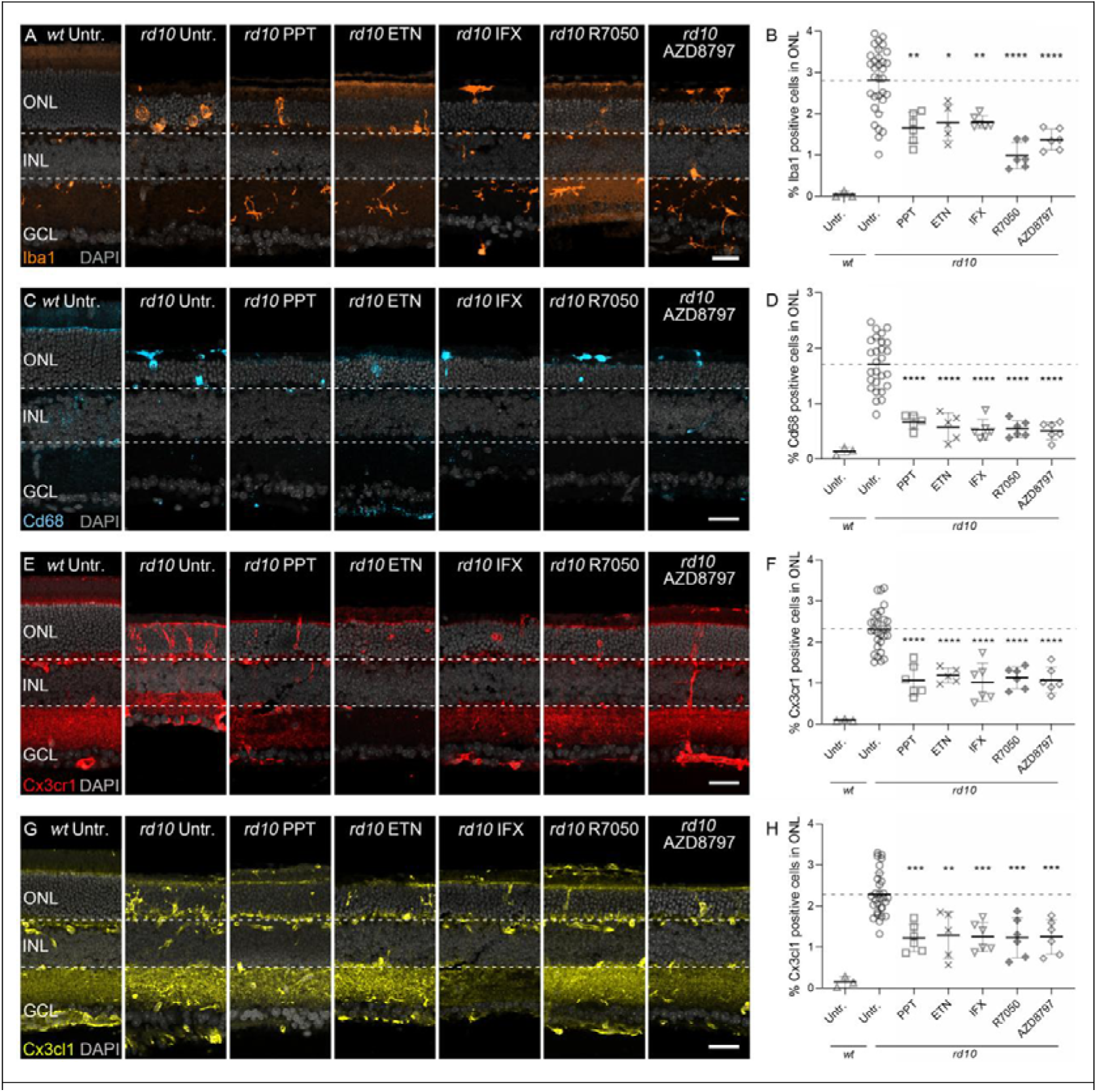
Esr1 activation, Tnf-α inhibition, and Cx3cr1 blockade suppress microglial recruitment and Cx3cl1/Cx3cr1 signaling in *rd10*. **(A–B)** Immunostaining of Iba1 revealed robust microglial recruitment into the ONL in *rd10* in vivo, which was significantly reduced by Esr1 activation (PPT), Tnf-α blockade (ETN, IFX), Tnfr1 antagonism (R7050), and Cx3cr1 inhibition (AZD8797). **(C-D)** Cd68 Immunostaining followed a similar trend: minimal in untreated *wt*, upregulated in *rd10*, and attenuated following intervention. **(E–H)** Immunostaining showed that Cx3cr1 (red) and Cx3cl1 (yellow) expression, barely detectable in *wt* retinas, were markedly increased in *rd10* and suppressed by all treatments. * = *p* < 0.05; ** = *p* < 0.01; *** = *p* < 0.001; **** = *p* < 0.0001. ONL = outer nuclear layer, INL = inner nuclear layer, GCL = ganglion cell layer, scale bar = 50 µm. Dashed lines indicate expression levels in untreated *rd10* controls. DAPI (grey) was used as nuclear counterstain. Statistical testing: one-way ANOVA with Tukey’s multiple comparison post hoc test.

Consistent with these changes, Cx3cl1 and Cx3cr1, barely detectable in untreated *wt*, were strongly upregulated in degenerating retinas but were consistently reduced by all interventions (Figure 6E-H; Figure S9C,D,G,H; Figure S10E-H). Notably, Tnf-α expression paralleled microglial and Cx3cl1/Cx3cr1 dynamics: ETN, IFX, R7050, and AZD8797 each produced significant and comparable suppression of Tnf-α across *rd1* explants, *in vivo rd10*, and ZAP-treated *wt* models (Figure S11A-F).

Together, these results indicate that Esr1 activation, Tnf-α/Tnfr1 inhibition, and Cx3cr1 blockade converge on a shared mechanism that dampens microglial recruitment, restrains Cx3cl1/Cx3cr1 signaling, and attenuates neuroinflammation across distinct degenerative contexts.

### Inflammatory signatures correlate with photoreceptor degeneration and are modulated by Esr1, Tnf-**α**, and Cx3cr1 inhibition

To enable cross-model comparisons, expression data were normalized by linear scaling, setting the minimum and maximum values to zero and one, respectively. In untreated *wt* retinas, both TUNEL positivity and inflammatory marker expression were consistently low relative to *rd1*, *rd10*, and ZAP-treated *wt* models (Figure D7A-C). Among inflammatory proteins, Tnf-α exhibited relatively elevated baseline expression even in *wt* retinas.

Pharmacological interventions targeting Esr1 (PPT), Tnf-α signaling (ETN, IFX, R7050), and Cx3cr1 (AZD8797) significantly reduced photoreceptor cell death and suppressed expression of Iba1, Cd68, Cx3cr1, Cx3cl1, and Tnf-α across all models. Despite this, residual Tnf-α levels remained comparatively higher in *rd10* and ZAP-treated *wt* retinas (Figure 7B-C), suggesting persistent or model-specific inflammatory resistance.

**Figure 7.**
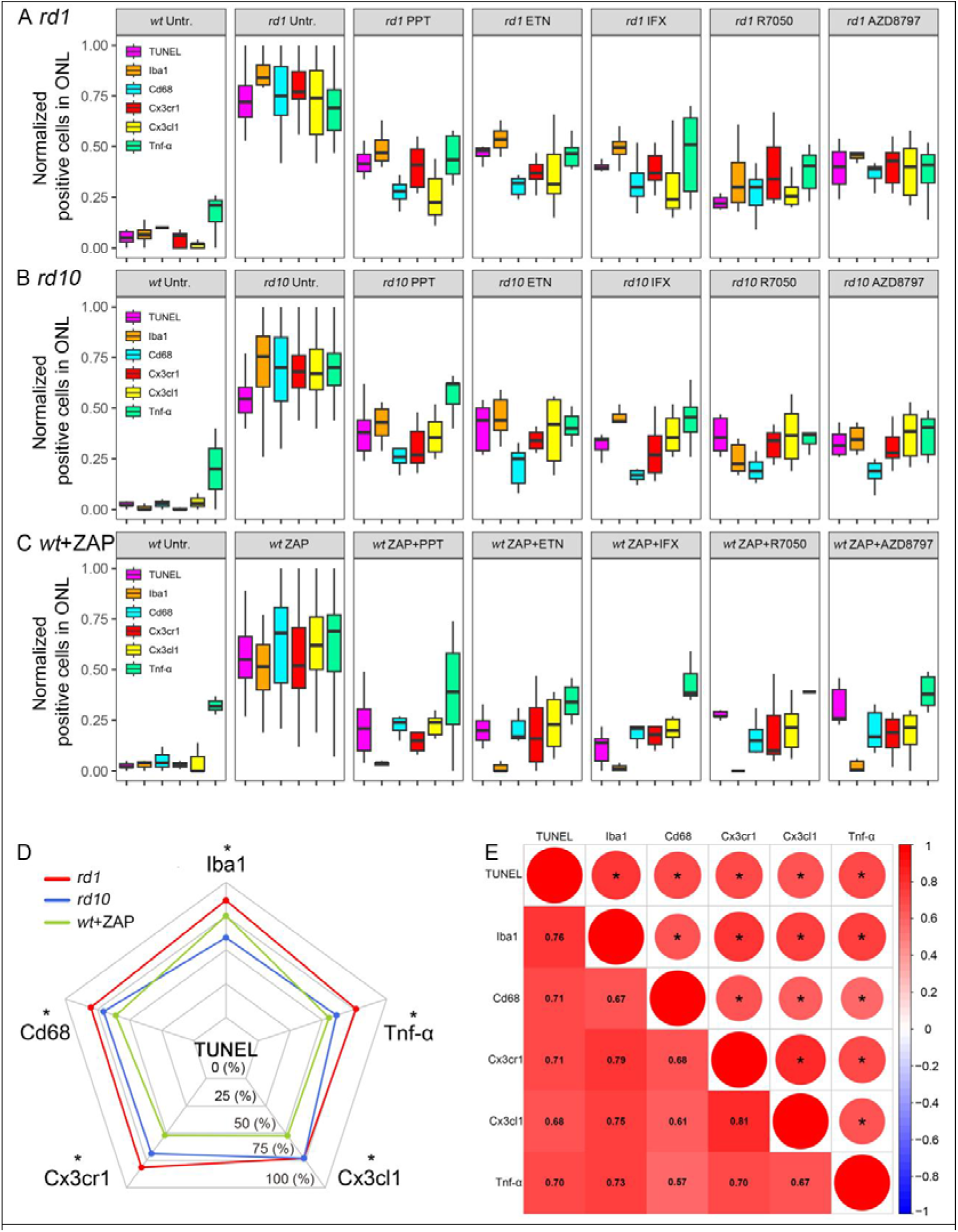
Inflammatory marker dynamics and their correlation with photoreceptor degeneration across models. **(A-C)** Comparison of Inflammatory markers across different experimental treatments in *rd1* (A), *rd10* (B), and *wt* subjected to ZAP-induced retinal degeneration (C). Normalized cell counts positive for TUNEL (magenta), Iba1 (orange), Cd68 (cyan), Cx3cr1 (red), Cx3cl1 (yellow), and Tnf-α (green) were shown across experimental groups. **(D)** Radar plots illustrating Spearman correlation between each inflammatory marker and TUNEL-positive cells in *rd1* (red), *rd10* (blue), and ZAP-treated *wt* (green) models. **(E)** Global Spearman correlation matrix among inflammatory markers and TUNEL across all degeneration models. Circle asterisks denote statistical significance; numeric values in squares indicate correlation coefficients (*R*).

Radar plot based Spearman analyses revealed strong positive correlations between photoreceptor death (TUNEL positivity) and inflammatory markers (Iba1, Cd68, Cx3cr1, Cx3cl1, Tnf-α) in all models examined (Figure 7D). A global Spearman analysis integrating data across *rd1*, *rd10*, and ZAP-treated *wt* further confirmed robust intercorrelations among these markers (Figure 7E), underscoring their co-regulation and central involvement in neuroinflammatory signaling.

Together, these findings implicate Tnf-α, Cx3cl1-Cx3cr1, and microglial markers (Cd68, Iba1) as tightly linked inflammatory mediators that collectively drive photoreceptor degeneration, and highlight their therapeutic modulation *via* Esr1, Tnf-α, or Cx3cr1 inhibition.

## Discussion

Our study establishes Esr1 as a pivotal neuroprotective modulator in *Pde6b*-related retinal degeneration, acting through suppression of microglial activation and downstream pro-inflammatory signaling. We observed robust upregulation of Esr1 in degenerating retinas, and selective pharmacological activation of Esr1 consistently mitigated photoreceptor loss, preserved dark-adapted ERG responses, and downregulated inflammatory mediators, including Tnf-α, Cx3cl1/Cx3cr1, Cd68, and Iba1, across *rd1*, *rd10*, and pharmacologically induced models. Conversely, blockade of Esr1 abolished these protective effects, underscoring its central role in maintaining photoreceptor survival. Interestingly, systemic administration of E2 produced divergent outcomes: neuroprotective in *rd1* yet deleterious in *rd10*, highlighting the genotype-dependent complexity of endogenous estrogen signaling. By contrast, selective Esr1 activation with PPT provided consistent, genotype-independent neuroprotection, offering a more predictable and clinically translatable strategy than global estrogen supplementation. Mechanistically, Esr1 activation transcriptionally repressed Tnf-α, curtailed Cx3cl1/Cx3cr1-driven microglial recruitment, and dampened microglial activation, thereby limiting retinal neuroinflammation and preserving photoreceptor integrity. Moreover, selective inhibition of Tnfr1 with R7050 significantly outperformed broad-spectrum Tnf-α inhibitors (ETN and IFX) in reducing ONL cell death and maintaining retinal function. Collectively, these findings identify Esr1 as an intrinsic protective mechanism in degenerating photoreceptors and nominate downstream effectors, particularly Tnfr1 and Cx3cr1, as actionable therapeutic targets for retinal neurodegeneration (Figure 8).

**Figure 8.**
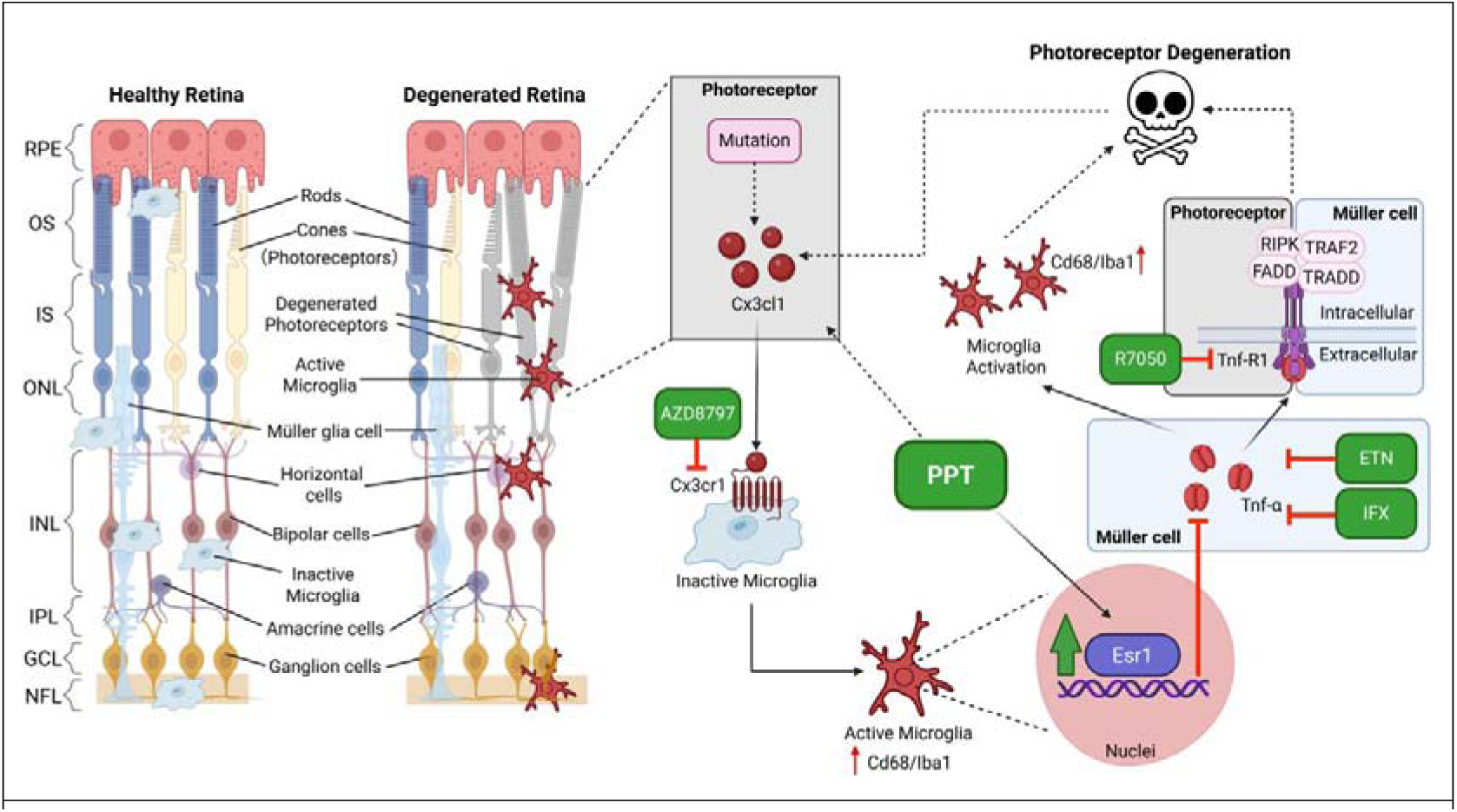
Schematic model of Esr1-mediated suppression of inflammatory signaling in *Pde6b*-associated retinal degeneration. In the healthy retina, microglia remain quiescent and are largely excluded from the outer nuclear layer (ONL). In *Pde6b*-mutant retinas, disease-associated stress induces upregulation of the chemokine Cx3cl1, which engages its receptor Cx3cr1 on microglia to drive their activation and migration into the ONL. Concomitantly, Esr1 expression is upregulated in degenerating retinas, suggesting a self-protective mechanism. Activated microglia transcriptionally upregulate Tnf-α and release it onto Müller cells, which in turn amplify microglial recruitment and activation. Simultaneously, Tnf-α engages Tnf receptors, particularly Tnfr1, on photoreceptors or Müller cells, initiating cell death cascades *via* recruitment of Tnfr1-associated death domain protein (TRADD) and subsequent assembly of a death-inducing signaling complex (DISC) comprising TRAF2, RIPK, and FADD. These signaling events promote direct and indirect photoreceptor degeneration while reinforcing a feed-forward loop that sustains Cx3cl1 expression and chronic neuroinflammation. Selective Esr1 activation by PPT interrupts this pathogenic cascade by suppressing Cx3cl1 expression in photoreceptors and repressing Tnf-α transcription in microglia, thereby disrupting microglia-driven neurotoxicity and preserving retinal integrity. The drugs used in this study and their targets are indicated. Red colour indicates destructive processes, while green labeled compounds promote photoreceptor survival.

### Diverse effects of E2 in *rd1* and *rd10* mouse

Estrogen is widely recognized for its neuroprotective effects. In our *rd1* retinal explant cultures, E2 reduced photoreceptor death, whereas in *rd10* explants it exacerbated degeneration, consistent with prior findings that E2 may induces retinal degeneration rather than protection [24, 25]. This opposing effect likely comes from the overactivation of estrogen receptor GPER1. *Gper* expression peaks at approximately P14 in mice [40]. Given that *rd1* explants were cultured from P5-P11 and *rd10* from P12-P20, the latter coincides with peak Gper1 activity. Overactivation of GPER1 may drive non-genomic signaling (cAMP/PKA, PI3K/Akt, MAPK) [41, 42], which enhances mitochondrial respiration and electron leakage, generating excessive reactive oxygen species (ROS) [43, 44]. Under conditions of glutathione depletion, iron accumulation, and sustained microglial activation, catechol estrogen-derived semiquinone/quinone intermediates undergo Fenton reactions, amplifying ROS [45, 46]. Concurrently, GPER1-driven Ca^2+^ influx aggravates glutamate excitotoxicity and mitochondrial dysfunction [47]. These convergent events probably shift E2 from neuroprotective in *rd1* to neurotoxic in *rd10*, accelerating photoreceptor degeneration.

### Esr1 as an intrinsic self-protective mechanism in degenerating retina

RNA-seq profiling of *rd1* and *rd10* retinas revealed positive enrichment of estrogen-related pathways, consistent with previous reports implicating endogenous estrogen as neuroprotective mechanisms in retinal stress responses [22]. However, the precise mechanisms underlying this self-protective pathway remain poorly defined. Althought the protective effects of estrogen have been described as genotype-dependent [24, 25],our data suggest that Esr1 may function as a more conserved axis of defense. scRNA-seq analyses showed that estrogen-responsive gene signatures were present in *wt* photoreceptor but absent in *Pde6b*-mutant counterparts, underscoring a loss of intrinsic estrogen responsiveness in the degenerating retina. Immunohistochemistry further demonstrated marked upregulation of Esr1 in *rd1* and *rd10* retinas, supporting the notion that Esr1 induction constitutes a compensatory response to photoreceptor stress. Importantly, while global estrogen signaling remains difficult to interpret due to its pleiotropic receptor interactions [20], Esr1 emerges as a more specific mediator. These findings suggest that Esr1 upregulation reflects an endogenous attempt by photoreceptors, Müller glia, or microglia to restrain inflammatory injury and preserve retinal integrity. Pharmacological reinforcement of this pathway amplifies this intrinsic protective program, shifting Esr1 from a compensatory mechanism to a therapeutically actionable axis. Thus, Esr1 activation can be conceptualized not merely as an exogenous intervention but as the potentiation of a built-in, self-protective defense against photoreceptor degeneration.

### The rescue effect of Esr1 activation likely arises from targeting retinal microglia

Unlike the variable effects observed with systemic E2, selective Esr1 blockade consistently exacerbated photoreceptor loss in both *rd1* and *rd10* explants, whereas Esr1 activation reproducibly attenuated cell death across all models, underscoring its robust neuroprotective effect. scRNA-seq analysis revealed that *Tnf* transcripts were confined to microglia, while immunostaining localized Tnf-α protein predominantly to Müller glia, supporting a paracrine model in which microglia-derived Tnf-α acts on Müller cells that express high levels of TNFR1 [48, 49]. Our immunohistochemical analyses confirmed Tnfr1 expression in Müller glia, consistent with their role as amplifiers of neuroinflammation through gliotic transformation, impaired glutamate clearance, and secondary release of chemokines such as Cx3cl1 [49, 50]. Although Esr1 staining was broadly detected across the retina, with enrichment in the IS, INL, and GCL, transcriptomic profiling following PPT treatment demonstrated significant repression of *Tnf* expression. This aligns with prior work showing that estrogen-bound Esr1 can tether NF-κB p65 and recruit corepressors to suppress *Tnf* transcription [51]. Consistently, PPT-mediated Esr1 activation markedly reduced Tnf-α protein levels in *rd1*, *rd10*, and ZAP-induced *wt* models, alongside parallel reductions in Cd68, Iba1, and Cx3cl1/Cx3cr1 expression.

Taken together, these findings suggest microglia are the principal source of Tnf-α in degenerating retinas, and Esr1 activation exerts its protective effects primarily by suppressing microglial Tnf-α transcription. While the contribution of Esr1 signaling in other retinal cell types cannot be excluded, our data strongly suggest that targeting microglial Esr1 is probably central to disrupting Tnf-α-driven neuroinflammation and preserving photoreceptor integrity.

### Inhibition of Tnf-**α** signaling, particularly Tnfr1, mitigates photoreceptor degeneration

TNF-α is a central mediator of neuroinflammation [52], predominantly produced by activated macrophages and microglia [53, 54]. It is initially synthesized as a type II transmembrane protein (tmTNF) and subsequently cleaved by TNF-α-converting enzyme (TACE/ADAM17) into its soluble form (sTNF) [55]. While sTNF selectively activates TNFR1, tmTNF can engage both TNFR1 and TNFR2 [53]. TNFR1 is strongly linked to cell death through the recruitment of TNFR1-associated death domain protein (TRADD), which assembles with TNFR-associated factor 2 (TRAF2), receptor-interacting protein kinase (RIPK), and Fas-associated death domain protein (FADD) to form the Death-Inducing Signaling Complex (DISC), initiating programmed cell death [56].

Our data revealed that Tnf-α expression remained elevated compared to other inflammatory markers (including Iba1, Cd68, Cx3cr1, and Cx3cl1), even in untreated *wt* retinas. This preferential upregulation of Tnf-α may be attributed to the photoreceptor outer segment (OS) phagocytosis [57]. Photoreceptor OS discs are constantly synthesized at the base and shed at the distal tip, a dynamic process critical for sustaining phototransduction efficiency and removing oxidatively damaged or dysfunctional components [57–59]. However, this high rate of metabolic and structural turnover may inherently generate low-grade, non-pathological inflammatory or stress signals [60], thereby contributing to baseline Tnf-α expression, even in the absence of overt degeneration.

In our *rd1*, *rd10*, and ZAP-induced degeneration models, Esr1 activation consistently suppressed Tnf-α expression, and pharmacological inhibition of Tnf-α signaling further confirmed its pathogenic role. ETN and IFX neutralized both sTNF and tmTNF [55], thereby broadly inhibiting TNFR1 and TNFR2 signaling [53]. In contrast, R7050, a selective TNFR1 antagonist [61], conferred the most pronounced protective effect on photoreceptor survival. This aligns with the notion that TNFR1, which contains a death domain, primarily mediates inflammatory and cytotoxic responses [53, 56]. Indeed, our immunostaining confirmed Tnfr1 expression also within photoreceptor inner segments, while scRNA-seq revealed that *Tnfrsf1a* is absent in *wt* rods but aberrantly induced in *Pde6b*-mutant rods, suggesting that Tnfr1 may directly trigger photoreceptor death. By contrast, TNFR2 could promote pro-survival signaling. Indeed, TNFR2-mediated Etk phosphorylation has been shown to partially activate VEGFR2 [62], contributing to cell survival and proliferation [62, 63]. Thus, the non-selective blockade of Tnfr2 by ETN and IFX may partially offset the neuroprotective benefits of Tnfr1 inhibition. Collectively, these results highlight Tnfr1 as the principal effector of Tnf-α-induced photoreceptor degeneration in *Pde6b*-deficient retinas.

### Microglial activation and Cx3cl1-Cx3cr1 signaling drive progressive photoreceptor degeneration

In *rd1*, *rd10*, and ZAP-induced models of retinal degeneration, neuronal stress triggers a sustained inflammatory response characterized by microglial activation. Among the key mediators, the CX3CL1-CX3CR1 signaling axis emerged as a central regulator [39]. Stress-induced activation of NF-κB and AP-1 promotes CX3CL1 transcription and accumulation of its membrane-bound precursor [64]. As pathology progresses, proteolytic cleavage by Ca² /ROS-activated ADAM10, MAPK/TNF-driven ADAM17 (TACE), and microglial cathepsin S releases soluble CX3CL1 [65–67]. Acting as a damage-associated molecular pattern (DAMP), soluble CX3CL1 engages microglial CX3CR1 to promote recruitment and activation within the photoreceptor [18]. While transient fractalkine signaling may facilitate debris clearance [68], sustained CX3CL1-CX3CR1 engagement reinforces microglial reactivity and drives chronic neuroinflammation [18, 19]. We observed persistent upregulation of microglial markers (Cd68, Iba1) and fractalkine axis components (Cx3cl1, Cx3cr1) in both *rd1* and *rd10* retinas, supporting the role of prolonged microglial activation in disease progression. Notably, pharmacological inhibition of Cx3cr1 with AZD8797 robustly reduced ONL cell death across models, confirming this pathway as a critical effector of microglia-mediated neurotoxicity and a viable target for therapeutic intervention.

## Conclusion

The role of estrogen signaling in IRDs has been debated due to its context-dependent and often opposing effects. Here, we identify Esr1 as a central immunoregulatory hub in *Pde6b*-associated retinal degeneration. Esr1 is robustly upregulated in degenerating retinas, likely representing an intrinsic self-protective response, and its selective pharmacological activation confers consistent, genotype-independent neuroprotection across *rd1*, *rd10*, and ZAP-induced degeneration models, whereas Esr1 inhibition abolishes these benefits. In contrast, systemic E2 treatment elicits divergent, genotype-specific outcomes, underscoring the complexity of global estrogen signaling and highlighting the therapeutic advantage of receptor-selective modulation. Mechanistically, Esr1 activation interrupts a self-sustaining Cx3cl1/Cx3cr1-Tnf-α inflammatory loop that drives microglial recruitment, activation, and ONL damage. Interventions at critical nodes, including Esr1 activation (PPT), Tnf-α neutralization (ETN, IFX), Tnfr1-specific inhibition (R7050), and Cx3cr1 blockade (AZD8797), consistently suppressed inflammatory mediators, dampened microglial toxicity, preserved retinal architecture, and improved visual function, with Tnfr1 inhibition yielding the strongest protection. These findings establish Esr1 activation not merely as an external intervention but as the amplification of an endogenous defense circuit in the degenerating retina, and support a rational therapeutic framework in which Esr1 and its downstream inflammatory effectors are targeted to suppress microglia-driven neurotoxicity and safeguard photoreceptor survival in IRDs and other retinal degenerations.

## ACKNOWLEDGEMENTS

The authors would like to thank Huiqin Li (from Yunnan Infectious Disease Hospital) for excellent assistance.

## Contributions

Conceptualization, J.Y., ZL.H., JC.Y. and F.P.-D.; methodology, J.Y., L.W., YT.L., YD.L. and JR.L.; software, QL.Y., X.Y., YJ.D. and J.Y.; validation, J.Y. and L.W.; formal analysis, QL.Y., Y.L., WJ.Z. and QX.Y.; investigation, J.Y., YT.L., YD.L., JR.L. and L.W; data curation, J.Y., ZL.H., KW.J. WR.X. and JC.Y.; writing-original draft preparation, J.Y. and QL.Y.; writing—review and editing, F.P-D., C.S., D.C. and ZZ.Z.; visualization, J.Y. and QL.Y.; supervision, J.Y., ZL.H. and JC.Y.; project administration, J.Y., ZL.H. and JC.Y.;; funding acquisition, J.Y., ZL.H. and JC.Y. All authors have read and agreed to the published version of the manuscript.

