## Supplemental materials 1 for "Resolving the estrogen paradox in hereditary retinal degeneration: Esr1 activation suppresses Tnf-α signaling as a photoreceptor self-protection mechanism"

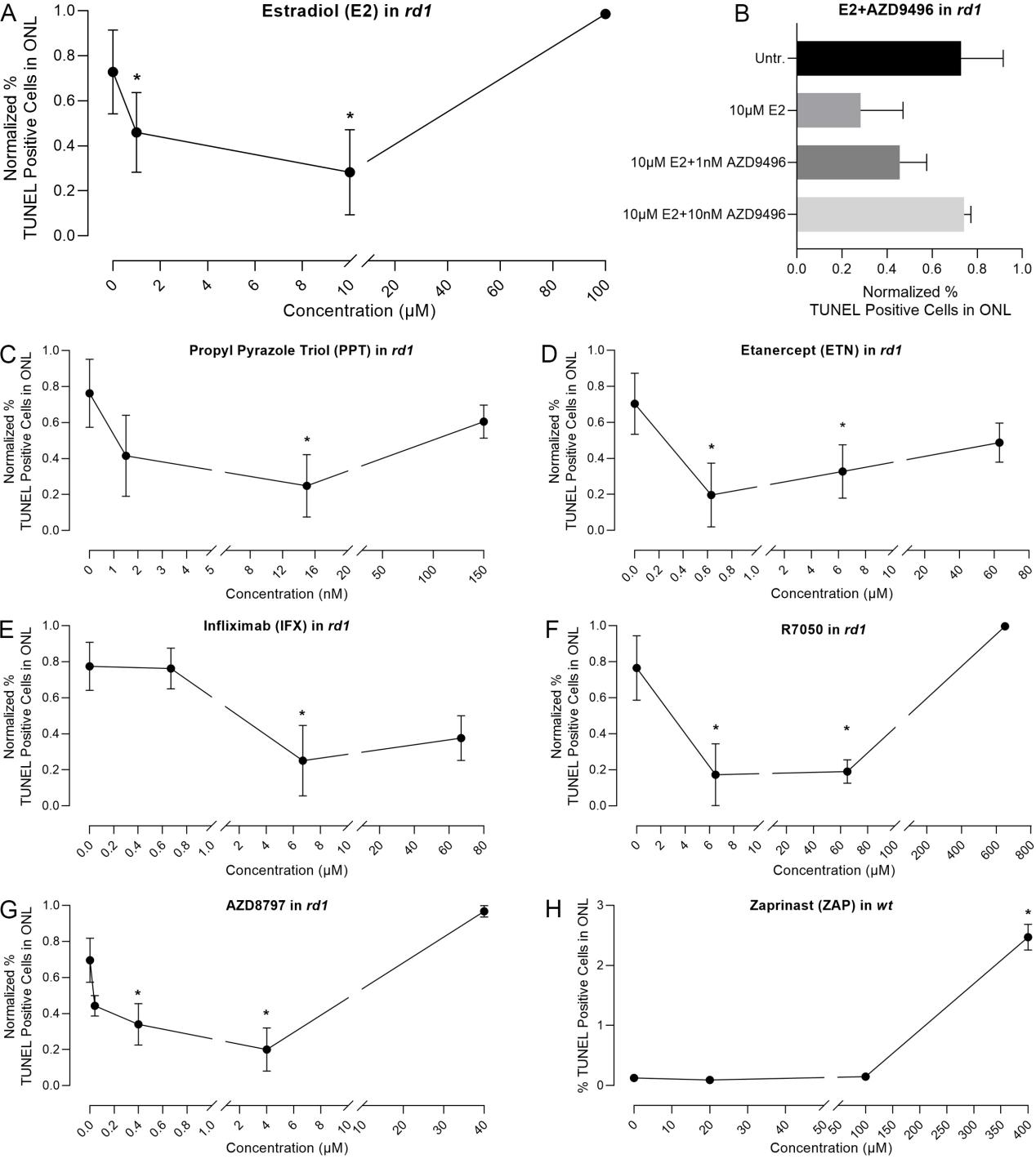

**Figure S1.** Dose-response curves for E2, E2+AZD9496, PPT, ETN, IFX, R7050, AZD8797 and ZAP. **(A)** Dose-response analysis of estradiol (E2) revealed significant neuroprotection at 1 and 10µM, with reduced TUNEL-positive cell death in the outer nuclear layer (ONL). **(B)** Co-administration of the Esr1 antagonist AZD9496 with E2 showed that 10nM AZD9496 fully abolished the protective effects of E2. **(C)** Propyl pyrazole triol (PPT), a selective Esr1 agonist, significantly reduced ONL cell death at 15nM. **(D-E)** Tnf-α neutralization using etanercept (ETN; 0.63 and 6.3µM) or infliximab (IFX; 6.7µM) significantly attenuated TUNEL-positive cell death in the ONL. **(F)** The Tnfr1-specific antagonist R7050 provided neuroprotection at 6.5 and 65µM. **(G)** Cx3cr1 inhibition with AZD8797 significantly reduced ONL cell death at 0.4 and 4µM. **(H)** Zaprinast (ZAP) induced significant photoreceptor toxicity only at 400µM in *wt* mice following intravitreal injection. Statistical significance was assessed using one-way ANOVA and Tukey’s multiple comparison post hoc test.

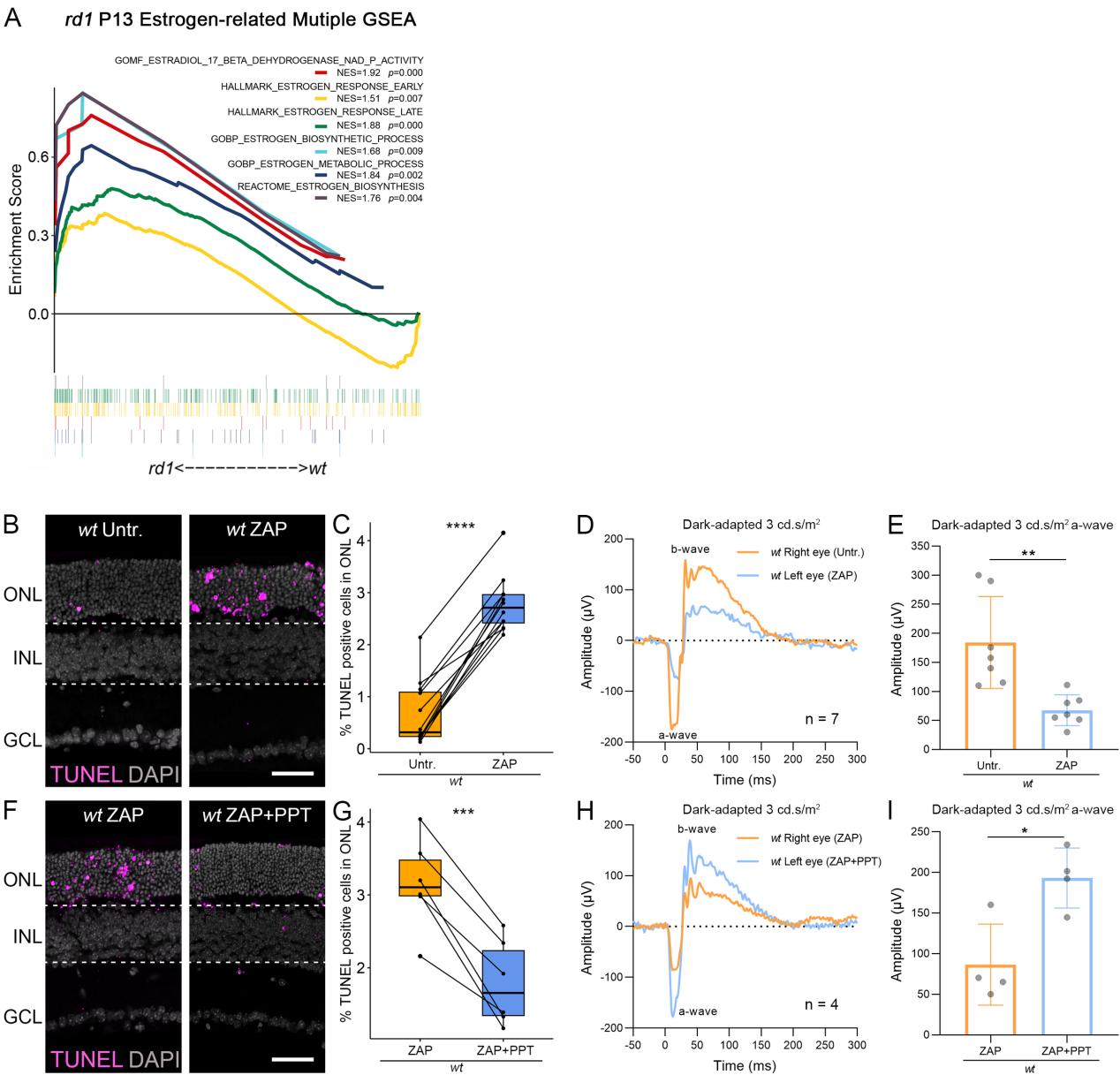

**Figure S2.** Estrogen signaling enrichment and Esr1 activation preserve photoreceptor integrity in ZAP-induced retinal degeneration. **(A)** GSEA of RNA-seq data from *rd1* retinas at postnatal day 13 (P13) revealed significant positive enrichment of Estrogen-related pathways.

**(B-E)** Zaprinast (ZAP, 400μM) in *wt* increased ONL cell death (B,C) and reduced ERG amplitudes (D,E) versus contralateral eyes. **(F-I)** Co-treatment with PPT (15nM) attenuated ZAP-induced ONL death (F,G) and rescued ERG responses (H,I). Error bars: SD; significance levels: * = *p* < 0.05; ** = *p* < 0.01; *** = *p* < 0.001; **** = *p* < 0.0001. ONL = outer nuclear layer, INL = inner nuclear layer, GCL = ganglion cell layer; scale bar = 50 µm. TUNEL staining (magenta) indicates dying cells, DAPI (grey) as nuclear counterstain.

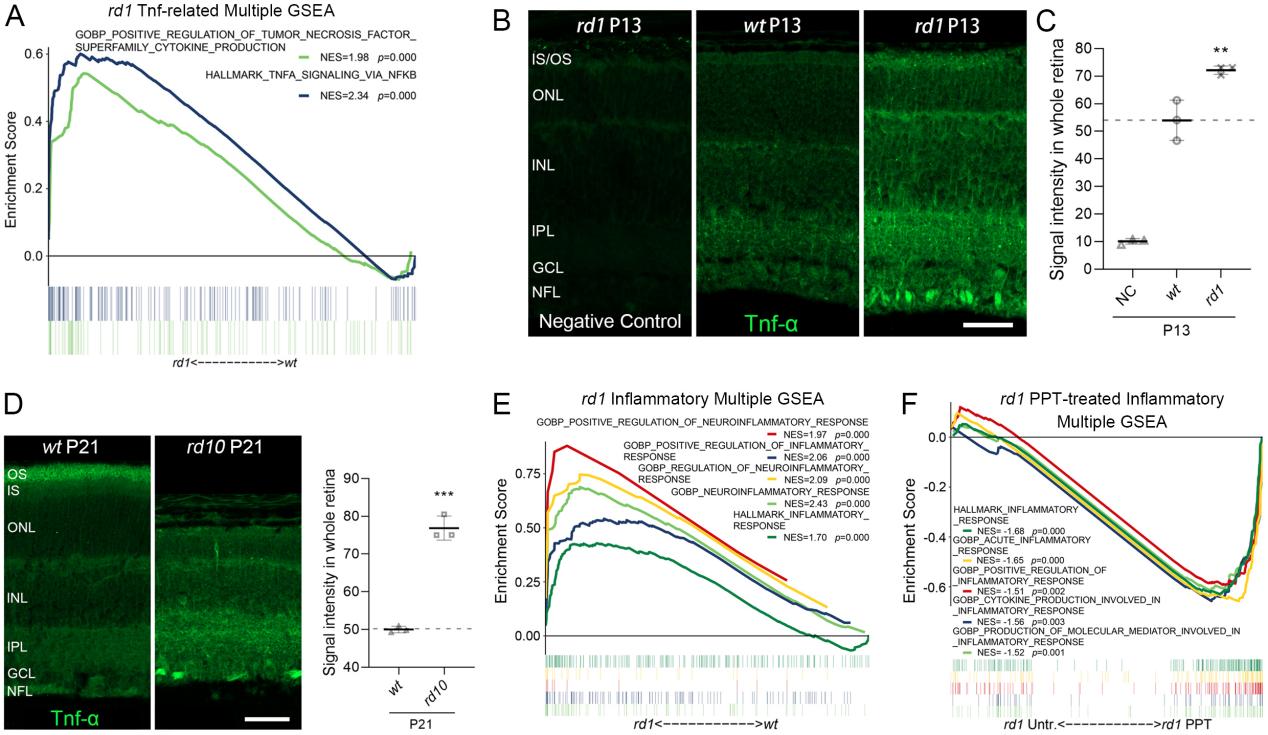

**Figure S3.** Neuroinflammation increased in *rd1* and *rd10*. **(A)** GSEA of RNA-seq data from *rd1* retinas at postnatal day 13 (P13) revealed significant positive enrichment of Tnf-related pathways. **(B-C)** Immunofluorescence staining for Tnf-α (green) demonstrated increased signal intensity in the entire retina of *rd1* mice compared to congenic wild-type (*wt*) controls and negative staining controls. **(D)** Tnf-α (green) expression was also elevated in *rd10* (P21) retinas relative to *wt*. **(E)** GSEA of *rd1* retinas at P13 similarly showed positive enrichment of inflammatory pathways. **(F)** GSEA of *rd1* retinal RNA-seq from explants showed enrichment of inflammatory associated pathways, reversed by PPT, DAPI (grey) as nuclear counterstain. Error bars: SD; significance levels: ** = *p* < 0.01; *** = *p* < 0.001. OS = outer segment, IS = inner segment, ONL = outer nuclear layer, INL = inner nuclear layer, IPL = inner plexiform layer, GCL = ganglion cell layer, NFL = nerve fiber layer; scale bar = 50µm. Dashed lines indicate expression levels in *wt* controls.

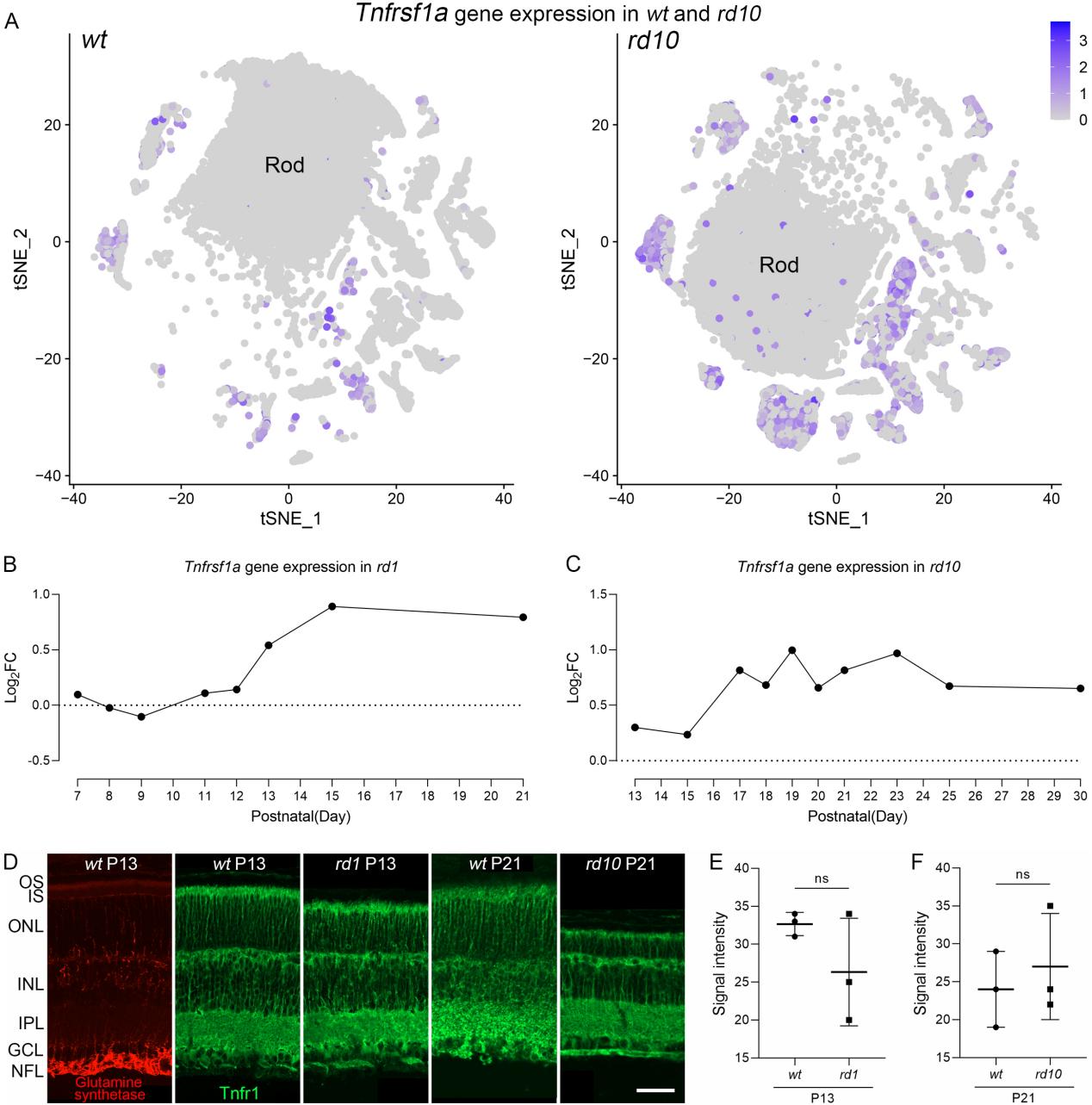

**Figure S4.** Tnfr1 gene and protein expression in degenerating retinas. **(A)** Feature plot revealed minimal expression of *Tnfrsf1a* in rod photoreceptors of wild-type (*wt*) retinas, whereas clear upregulation was observed in *rd10* rods. **(B-C)** Bulk RNA-seq analysis of *Tnfrsf1a* expression revealed a progressive upregulation from P7 to P21 in *rd1* (B) and from P13 to P30 in *rd10* (C), indicating increased transcription during disease progression. **(D)** Immunofluorescence staining for glutamine synthetase (GS; red) and Tnfr1 (green) indicated that Tnfr1 protein localized to the photoreceptor OS, Müller glial cells and INL. **(E-F)** Quantification of Tnfr1 protein signal intensity showed no significant difference between *rd1*, *rd10*, and their congenic *wt* controls. Error bars: SD; significance levels: ns = *p* > 0.01. OS = outer segment, IS = inner segment, ONL = outer nuclear layer, INL = inner nuclear layer, IPL = inner plexiform layer, GCL = ganglion cell layer, NFL = nerve fiber layer; scale bar = 50µm.

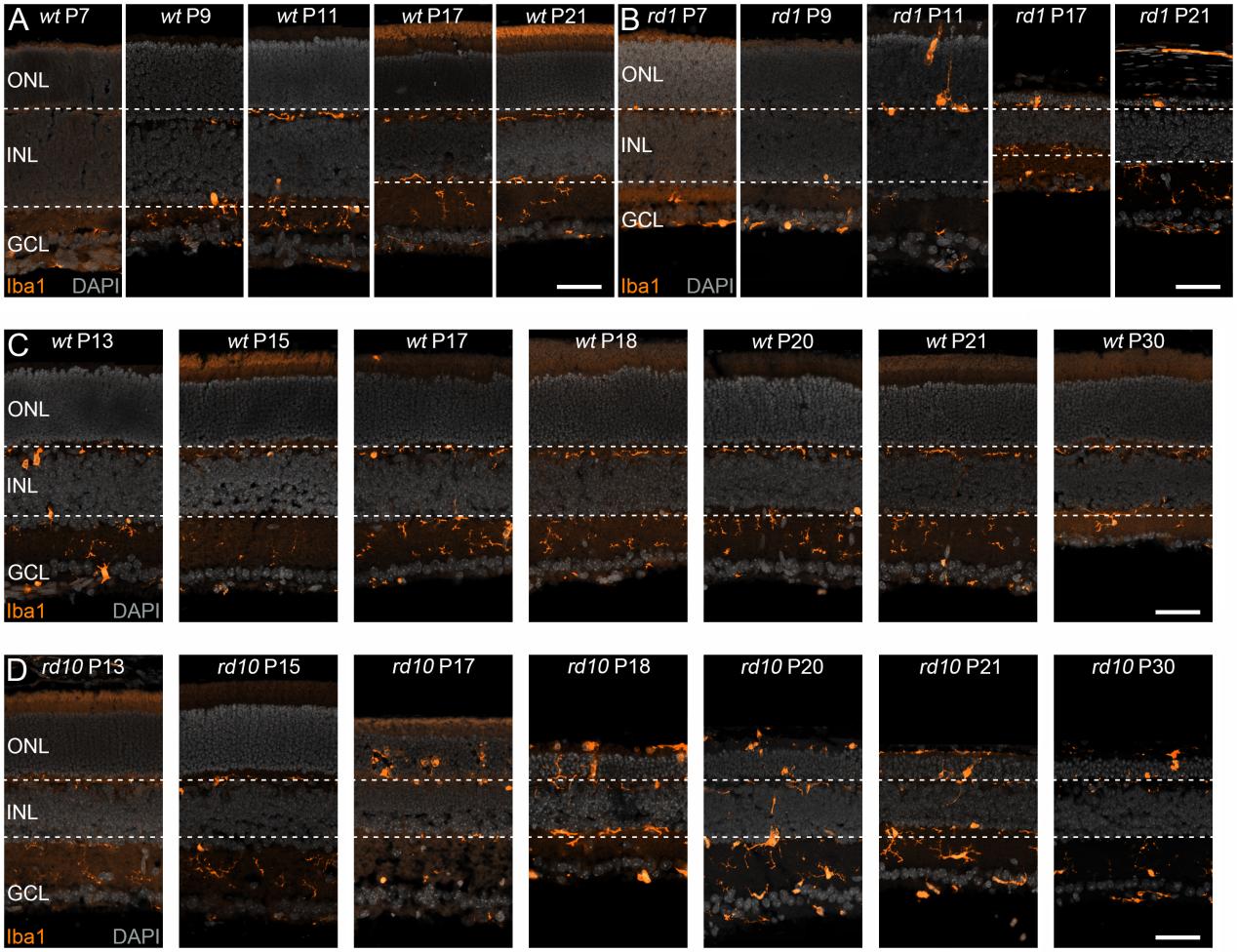

**Figure S5.** Temporal expression of Iba1 during retinal degeneration in *rd1*, *rd10*, and their congenic *wt* controls. **(A)** Immunostaining for Iba1 (orange) in *wt* retinas at postnatal days (P) 7, 9, 17, and 21 shows baseline microglial distribution. **(B)** In *rd1* retinas, Iba1 expression is absent at P7 and P9, but markedly increased at P17 and P21, indicating robust microglial activation and infiltration into ONL during degeneration. **(C)** Iba1 expression in *wt* retinas at P13, P17, P18, P20, P21, and P30 remains restricted to inner layers. **(D)** In *rd10* retinas, Iba1 signal is absent at P13, P15 and progressively increases from P17 to P30, with microglia migrating into the ONL, consistent with ongoing degeneration. DAPI (grey) was used as nuclear counterstain; ONL = outer nuclear layer, INL = inner nuclear layer, GCL = ganglion cell layer, scale bar = 50µm.

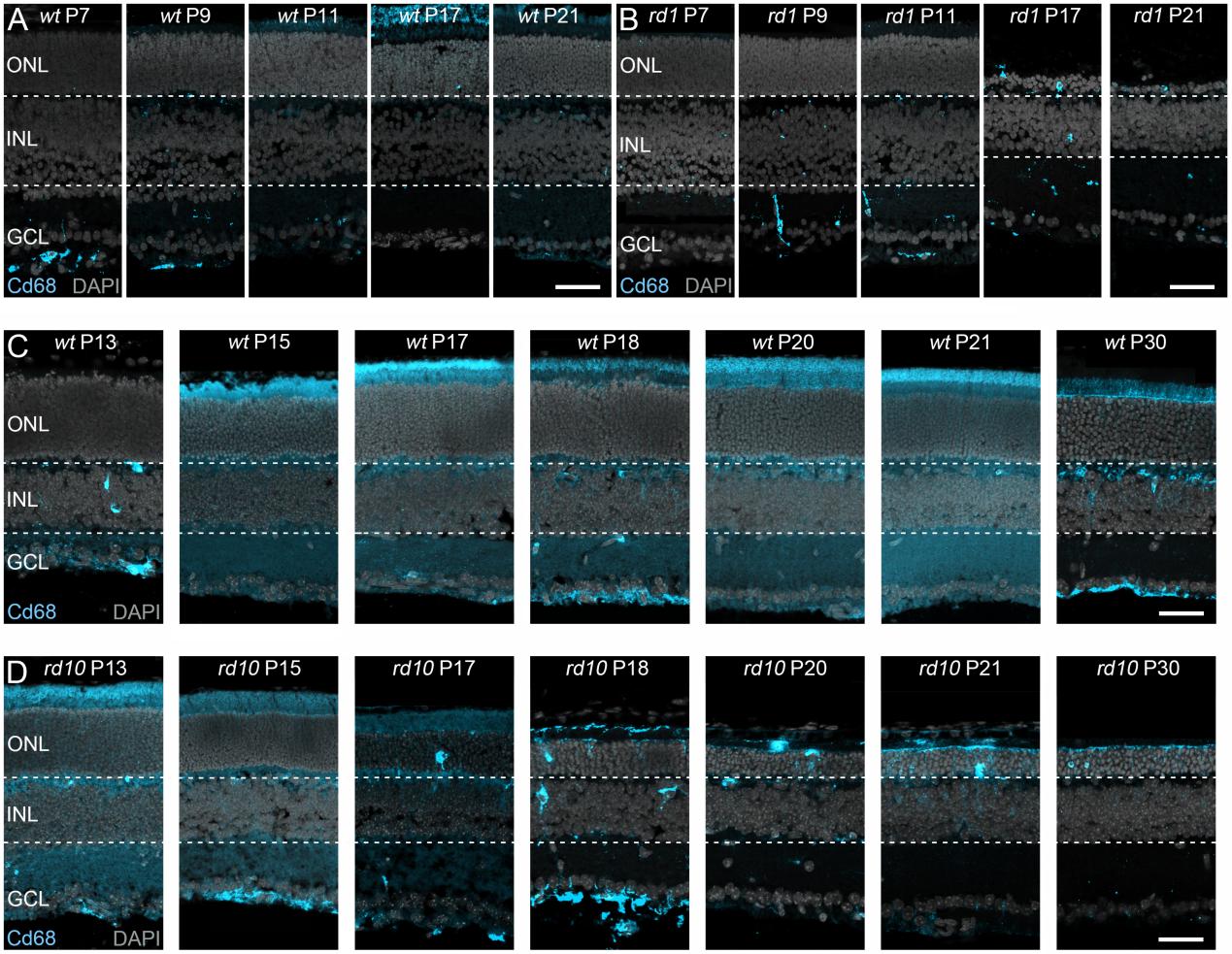

**Figure S6.** Temporal expression of Cd68 during retinal degeneration in *rd1*, *rd10*, and their congenic *wt* controls. **(A)** Immunostaining for Cd68 (cyan) in *wt* retinas at postnatal days (P) 7, 9, 17, and 21 shows minimal microglial activation. **(B)** In *rd1* retinas, Cd68 signal is absent at P7, P9, and P11 but becomes strongly elevated at P17 and P21, indicating robust microglial activation and infiltration into ONL during degeneration. **(C)** Cd68 staining in *wt* retinas at P13, P17, P18, P20, P21, and P30 remains confined to inner retinal layers. **(D)** In *rd10* retinas, while Cd68 staining is low at P13 and P15, it progressively increases at later time points, reflecting sustained microglial activation and accumulation within the ONL throughout degeneration. DAPI (grey) was used as nuclear counterstain; ONL = outer nuclear layer, INL = inner nuclear layer, GCL = ganglion cell layer, scale bar = 50µm.

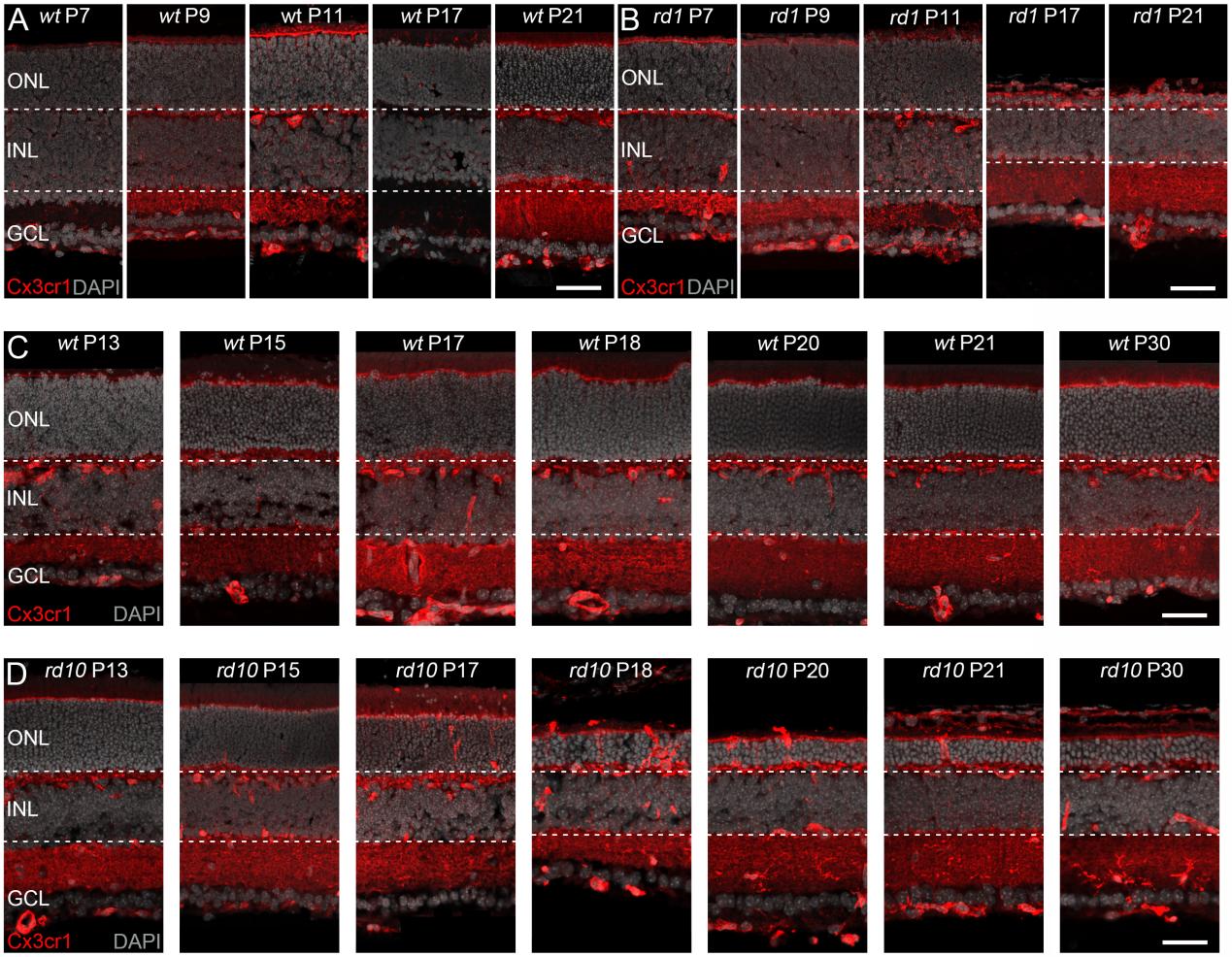

**Figure S7.** Temporal expression of Cx3cr1 during retinal degeneration in *rd1*, *rd10*, and their congenic *wt* controls. **(A)** Immunostaining for Cx3cr1 (red) in *wt* retinas at postnatal days (P) 7, 9, 11, 17, and 21 shows minimal protein expression. **(B)** In *rd1* retinas, Cx3cr1 positive cells are absent at P7, P9, and P11 but becomes strongly elevated at P17 and P21, indicating degeneration-associated expression. **(C)** In *wt* retinas, Cx3cr1 expression at P13, P15, P17, P18, P20, P21, and P30 remains confined to the inner retinal layers without ONL infiltration. **(D)** In *rd10* retinas, while Cx3cr1 staining is low at P13, and P15, it progressively increases at later time points, consistent with sustained retinal degeneration and inflammatory activation. DAPI (grey) was used as nuclear counterstain; ONL = outer nuclear layer, INL = inner nuclear layer, GCL = ganglion cell layer, scale bar = 50µm.

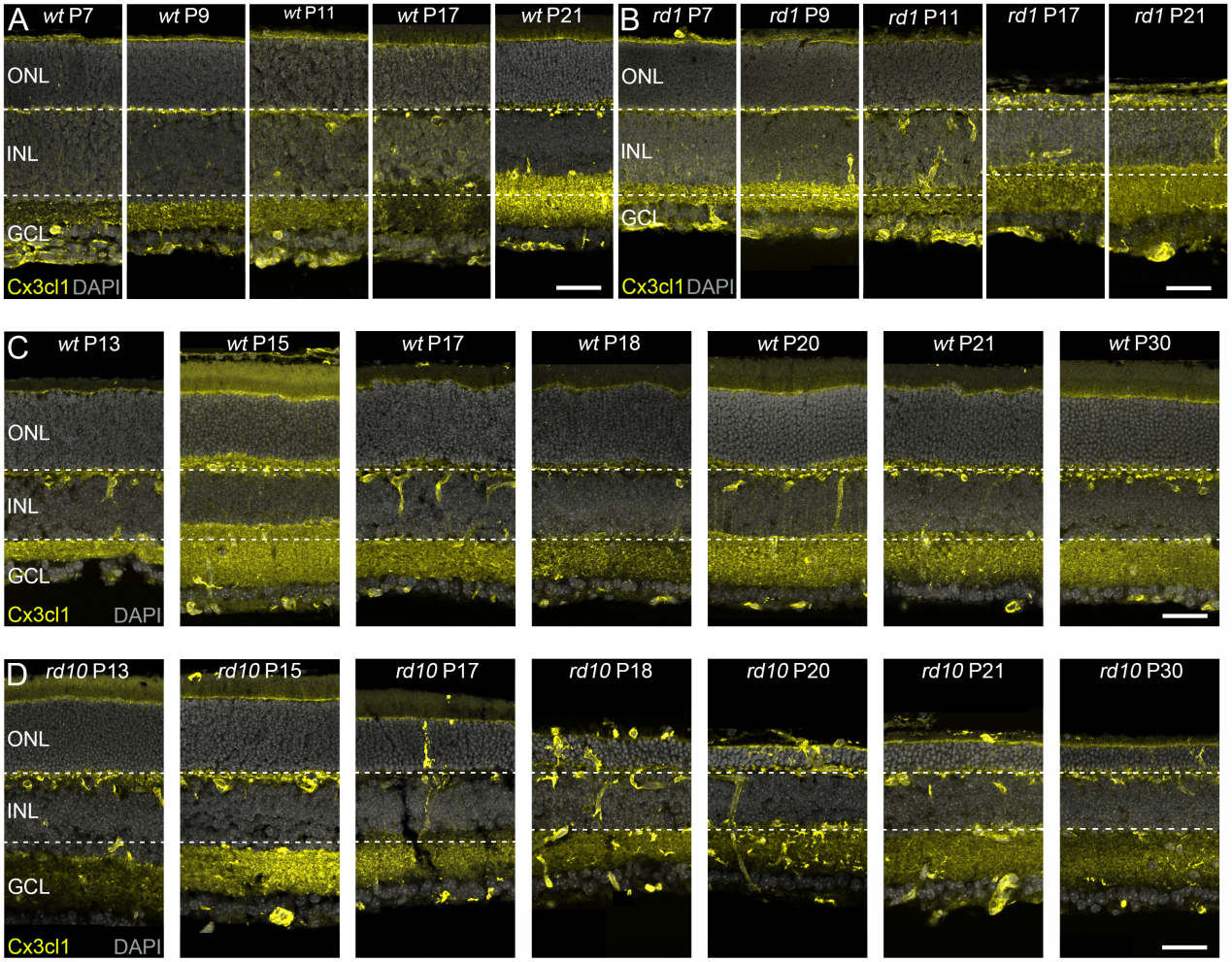

**Figure S8.** Temporal expression of Cx3cl1 during retinal degeneration in *rd1*, *rd10*, and their congenic *wt* controls. **(A)** Immunostaining for Cx3cl1 (yellow) in *wt* retinas at postnatal days (P) 7, 9, 11, 17, and 21 shows minimal protein expression. **(B)** In *rd1* retinas, Cx3cl1 positive cells are absent at P7, P9, and P11 but becomes strongly elevated at P17 and P21, indicating degeneration-associated expression. **(C)** In *wt* retinas, Cx3cl1 expression at P13, P15, P17, P18, P20, P21, and P30 remains confined to the inner retinal layers without ONL infiltration. **(D)** In *rd10* retinas, while Cx3cl1 staining is low at P13, and P15, it progressively increases at later time points, consistent with sustained retinal degeneration and inflammatory activation. DAPI (grey) was used as nuclear counterstain; ONL = outer nuclear layer, INL = inner nuclear layer, GCL = ganglion cell layer, scale bar = 50µm.

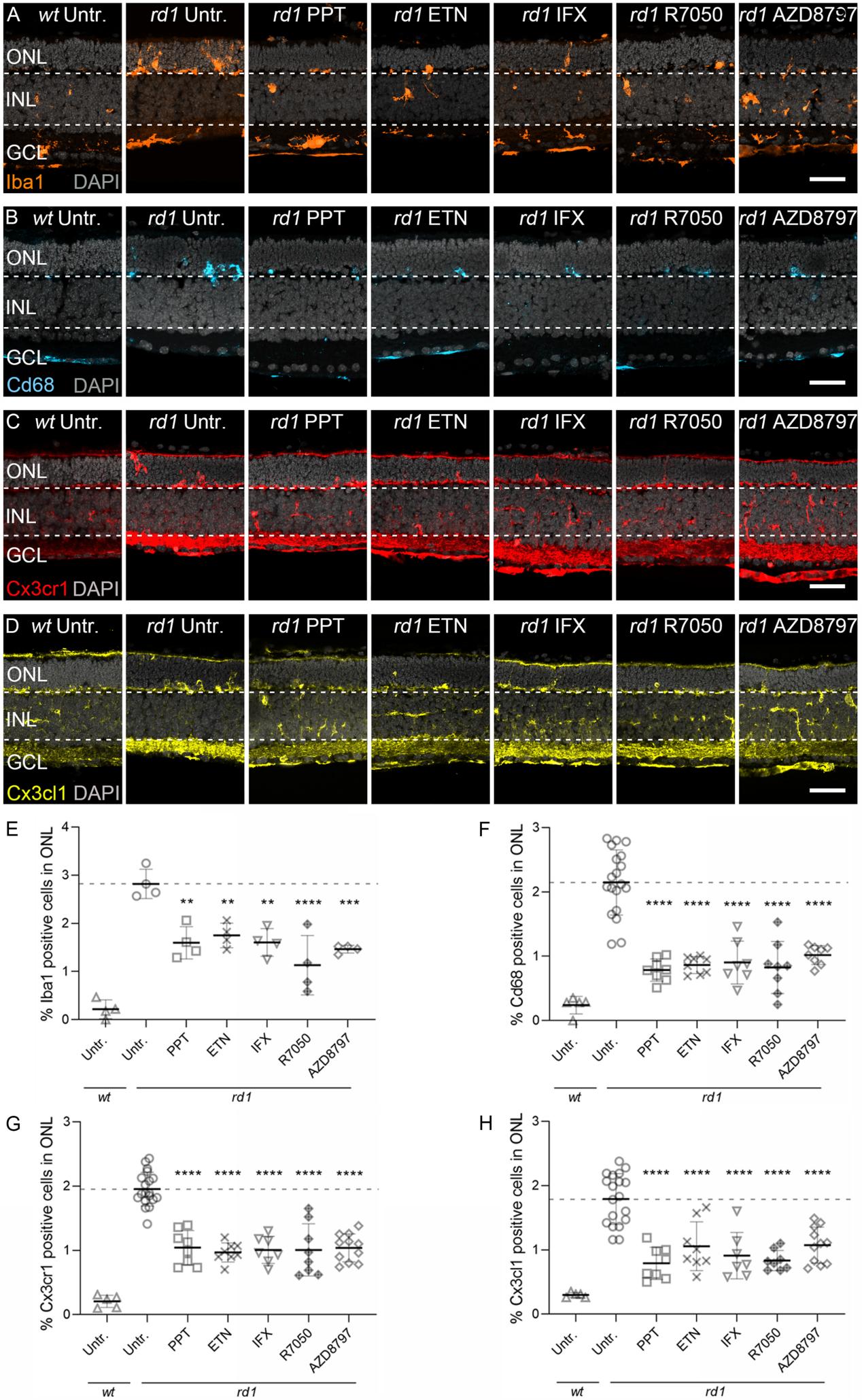

**Figure S9.** Esr1 activation, Tnf-α inhibition, and Cx3cr1 blockade suppress microglial recruitment and Cx3cl1/Cx3cr1 signaling in *rd1*. **(A, E)** Immunostaining of Iba1 revealed robust microglial recruitment into the ONL in *rd1* explants, which was significantly reduced by Esr1 activation (PPT), Tnf-α blockade (ETN, IFX), Tnfr1 antagonism (R7050), and Cx3cr1 inhibition (AZD8797). **(B, F)** Cd68 Immunostaining followed a similar trend: minimal in untreated *wt*, upregulated in *rd1*, and attenuated following intervention. **(C-D, G-H)** Immunostaining showed that Cx3cr1 (red) and Cx3cl1 (yellow) expression, barely detectable in *wt* retinas, were markedly increased in *rd1* and suppressed by all treatments. ** = *p* < 0.01; *** = *p* < 0.001; **** = *p* < 0.0001. ONL = outer nuclear layer, INL = inner nuclear layer, GCL = ganglion cell layer, scale bar = 50 µm. Dashed lines indicate expression levels in untreated *rd1* controls. DAPI (grey) was used as nuclear counterstain. Statistical testing: one-way ANOVA with Tukey’s multiple comparison post hoc test.

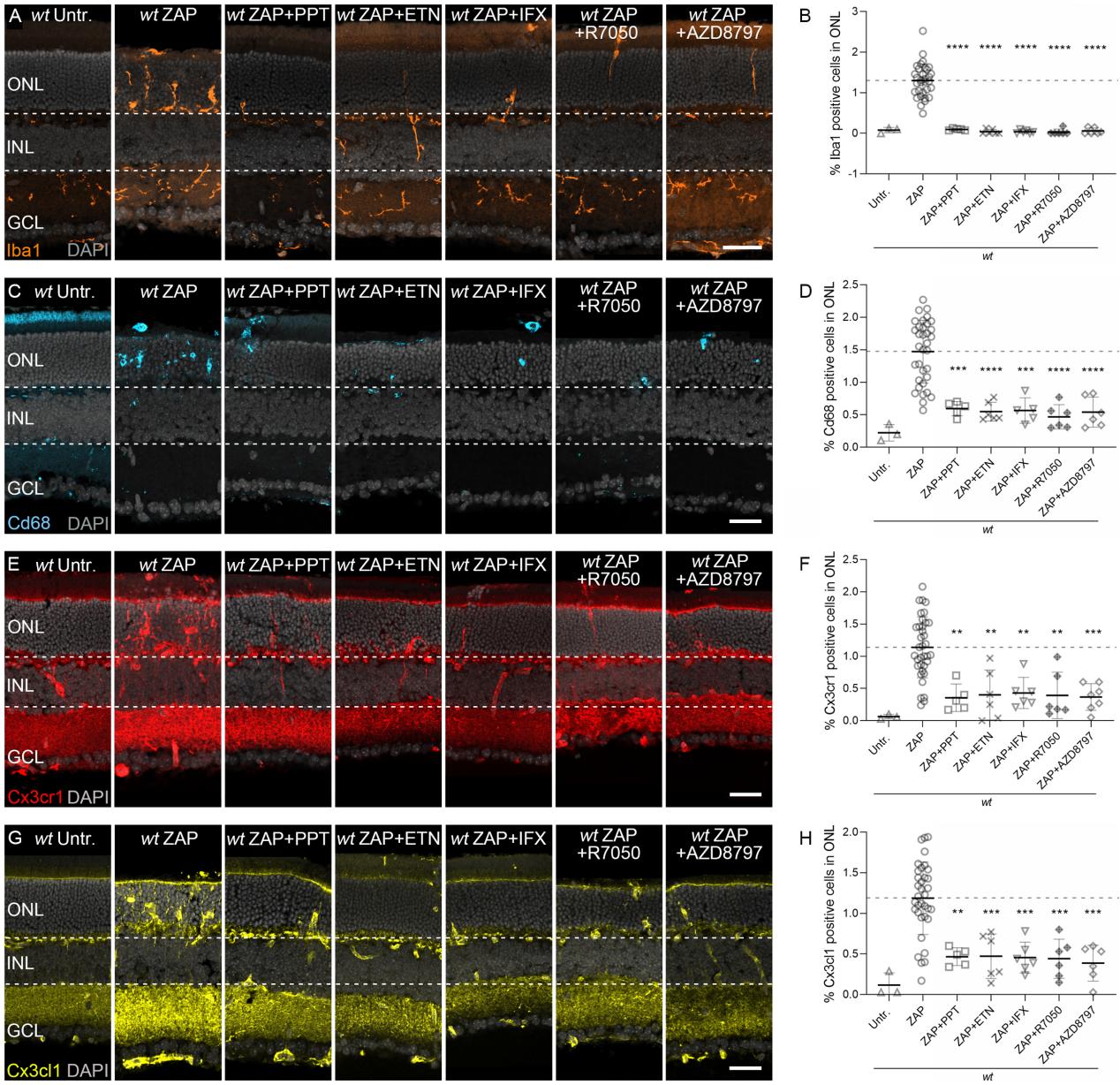

**Figure S10.** Esr1 activation, Tnf-α inhibition, and Cx3cr1 blockade suppress microglial recruitment and Cx3cl1/Cx3cr1 signaling in *wt* with ZAP-induced retinal degeneration. **(A, E)** Immunostaining of Iba1 revealed robust microglial recruitment into the ONL in *wt* with ZAP-induced retinal degeneration, which was significantly reduced by Esr1 activation (PPT), Tnf-α blockade (ETN, IFX), Tnfr1 antagonism (R7050), and Cx3cr1 inhibition (AZD8797). **(B, F)** Cd68 Immunostaining followed a similar trend: minimal in untreated *wt*, upregulated in *wt* with ZAP-induced retinal degeneration, and attenuated following intervention. **(E–H)** Immunostaining showed that Cx3cr1 (red) and Cx3cl1 (yellow) expression, barely detectable in untreated *wt* retinas, were markedly increased in *wt* with ZAP-induced retinal degeneration and suppressed by all treatments. ** = *p* < 0.01; *** = *p* < 0.001; **** = *p* < 0.0001. ONL = outer nuclear layer, INL = inner nuclear layer, GCL = ganglion cell layer, scale bar = 50 µm. Dashed lines indicate expression levels in ZAP-induced *wt*. DAPI (grey) was used as nuclear counterstain. Statistical testing: one-way ANOVA with Tukey’s multiple comparison post hoc test.

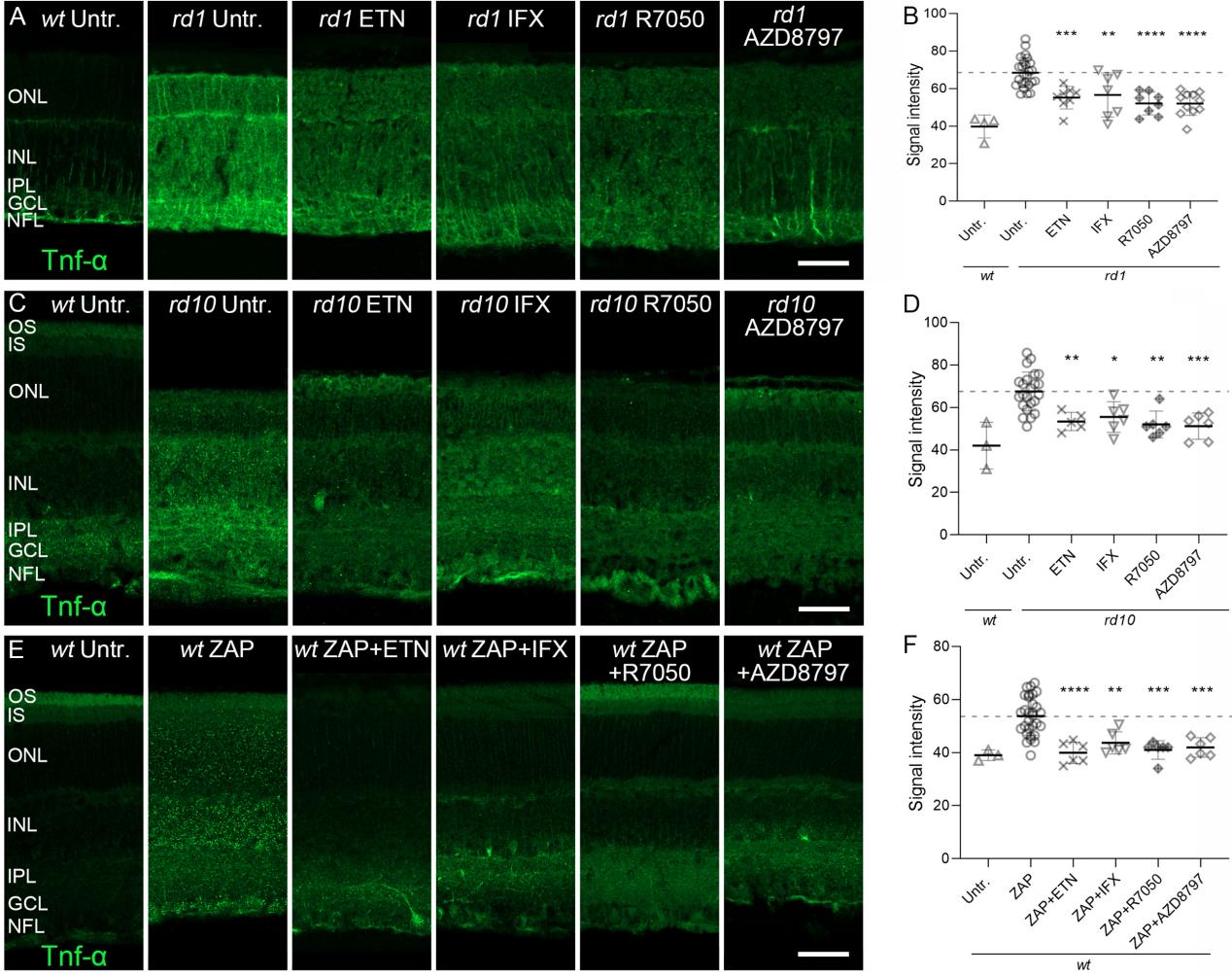

**Figure S11.** Tnf-α and Cx3cr1 inhibition suppress Tnf-α expression across RP models. **(A-B)** In *rd1* retinal explants, Tnf-α expression (green) was markedly elevated compared to untreated wild-type (wt) controls. Treatment ETN, and IFX (Tnf-α inhibitors), R7050 (Tnfr1 antagonist), or AZD8797 (Cx3cr1 inhibitor) significantly reduced Tnf-α signal intensity. **(C-D)** Similarly, *rd10* retinas exhibited widespread upregulation of Tnf-α, which was significantly suppressed following intravitreal administration of ETN, IFX, R7050, or AZD8797. **(E-F)** In *wt* mice subjected to ZAP-induced degeneration, Tnf-α expression was elevated throughout the retina. Co-treatment with ETN, IFX, R7050, or AZD8797 effectively reduced Tnf-α expression to near-baseline levels. Error bars: SD; significance levels: * = *p* < 0.05; ** = *p* < 0.01; *** = *p* < 0.001; **** = *p* < 0.0001. OS = outer segment, IS = inner segment, ONL = outer nuclear layer, INL = inner nuclear layer, IPL = inner plexiform layer, GCL = ganglion cell layer, NFL = nerve fiber layer; scale bar = 50 µm. Dashed lines indicate untreated or *wt* with ZAP-treated levels. Statistical testing: one-way ANOVA with Tukey’s multiple comparison post hoc test
